# Highly adaptable deep-learning platform for automated detection and analysis of vesicle exocytosis

**DOI:** 10.1101/2024.08.02.606323

**Authors:** Abed Alrahman Chouaib, Hsin-Fang Chang, Omnia M. Khamis, Santiago Echeverry, Lucie Demeersseman, Sofia Elizarova, James A Daniel, Salvatore Valitutti, Sebastian Barg, Constantin Pape, Ali H. Shaib, Ute Becherer

## Abstract

Vesicle exocytosis is a fundamental component of intercellular communication, in all organisms. It has been studied for decades, using various imaging tools. Nevertheless, exocytosis research is still limited by the lack of reliable automated analysis procedures. To address this, we developed the Intelligent Vesicle Exocytosis Analysis Platform (IVEA), a nearly universal solution for analyzing exocytosis acquired with live cell imaging. IVEA is applicable to a wide variety of experimental model systems, microscopes and reporter fluorophores. IVEA combines state-of-the-art deep-learning and computer vision regimes to enable fully automated analysis of large data. IVEA runs as a FIJI plugin and does not require prior training or human intervention. IVEA is 60 times faster than manual analysis and is able to detect rare events often missed by the human eye. Overall, IVEA represents a breakthrough in the analysis of cellular secretory mechanisms and has a transformative potential for the exocytosis imaging field.

## Introduction

Regulated exocytosis is a central process by which cells secrete substances on demand. It underlay synaptic transmission, hormone and cytokine release but also targeted release of cytotoxic protein by immune cells^1, 2^. It corresponds to a calcium triggered fusion of a vesicle with the plasma membrane leading to the release of biologically active substances. Understanding the underlying molecular mechanism is essential to develop therapies for mental disorders^3-5^, diabetes^6^ or immunological deficiencies such hemophagocytic lymphohistiocytosis^7^. Exocytosis can be monitored with live imaging techniques after over-expressing vesicular proteins tagged with fluorescent markers. Bulk vesicle exocytosis occurring during neuronal synaptic transmission, is visualized with pH sensitive fluorophores like the super ecliptic pHluorin (SEP) (**Fig. 1**) using various imaging techniques (confocal microscopy, epifluorescence microscopy, etc.)^8-10^. The imaging method of choice to observe individual vesicle before and during exocytosis in diverse cell types such as endocrine or immune cells with high temporal and spatial resolution is total internal reflection fluorescence microscopy (TIRFM)^11-13^. Combining TIRFM and pH dependent fluorophores enable the analysis of intermediate stages during individual vesicle fusion^10^. These techniques have resulted in a steadily rising volume of findings, but data analysis have emerged as a critical bottleneck that significantly hinders the production of large datasets. This challenge has become increasingly severe as scientific grade camera chips have moved to CMOS technology. As a result, the chips have grown in size, allowing for the capture of exceedingly expansive fields of view in which numerous cells or hundreds of synapses can be measured simultaneously. As the sensitivity of the camera increases, smaller pixel size can be used increasing the image resolution but also image size. Finally, the chip readout time has decreased allowing increased acquisition speeds resulting in the generation of vast quantity (terabytes) of image data. To mitigate this analysis, bottleneck several automatic methods have been developed^14-20^. However, the impact of these published methods has been limited due to either their complexity or lack of versatility for a diverse range of data sets.

**Figure 1.**
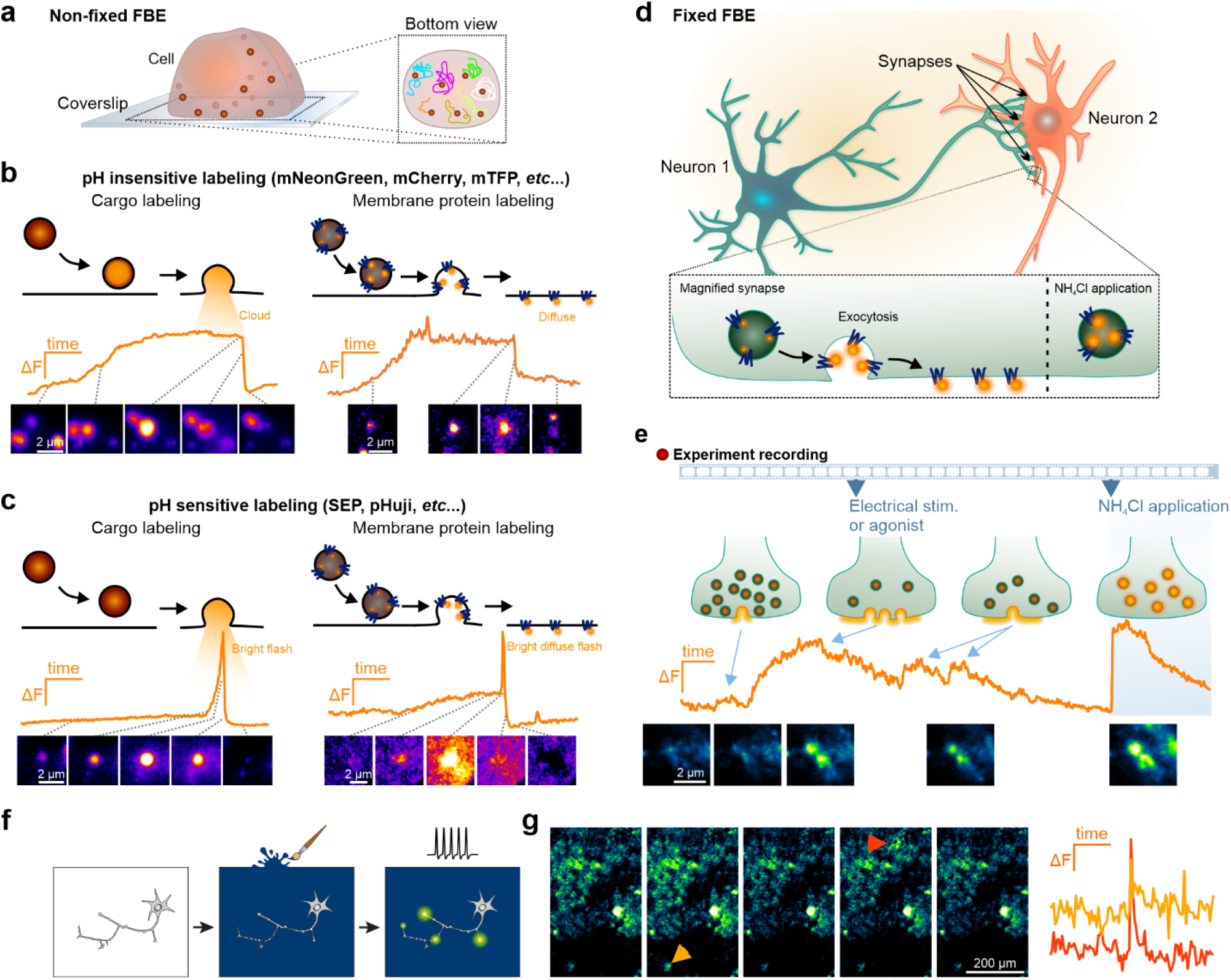
Vesicle exocytosis characteristics across different systems. **a**, Schematic representation of a T-cell in which the secretory vesicles, i.e. lytic granules, are labeled by a fluorescent marker. They are distributed in the entire cell (left) and move over time (right). **b-c**, Top row: schematic representation of the successive events occurring during exocytosis in T-cells. Middle: fluorescence intensity over time of a single vesicle shown below. Bottom: snapshots of the vesicle during exocytosis. The time points of the snapshots are indicated by a stippled line on the graph above. **b**, Examples of the exocytosis of vesicles labelled by pH insensitive fluorescent proteins that are either bound to a cargo protein (right, granzyme B) or a membrane protein (left, synaptobrevin2). **c**, Examples of the exocytosis of vesicles labelled by pH sensitive fluorescent proteins such as the super ecliptic pHluorin (SEP) that are either bound to a cargo protein (right, granzyme B) or a membrane protein (left, synaptobrevin2). Note that with the pH sensitive probe, fusion of the vesicle is accompanied by a sharp increase in fluorescence intensity due to neutralization of the intravesicular lumen. **d**, Scheme of synaptic transmission measured with synaptophysin-SEP (SypHy). The pH-sensitive fluorophore SEP is located in the acidic environment of the vesicle, which quenches its fluorescence. Upon exocytosis, SEP is exposed to the neutral pH (7.4) of the extracellular medium lightening it up, while NH4^+^-containing medium neutralizes the lumen of all the vesicles making them visible. **e**, Exemplary measurement of synaptic transmission with SypHy. Top: scenarios that may occur at individual synapses with their specific fluorescent intensity variation over time (Middle). The lower row exhibits snapshots of this synapse at the time points of the scenarios shown in the upper row. During non-synchronous exocytosis, small and brief increase in fluorescence is observed when very few vesicles fuse with the plasma membrane. Electrical or agonist stimulation causes simultaneous exocytosis of many vesicles, leading to a durable rise in fluorescence while SypHy remains in the plasma membrane. The fluorescence intensity decreases upon the endocytosis of SypHy and subsequent reacidification of the vesicle. NH4^+^ induces maximum fluorescence intensity increase. Note that changes in fluorescence intensity occur at fixed small locations, i.e. the synapse. **f**, Illustration of a dopaminergic neuron culture surrounded by a ‘painted’ fluorescent dopamine nanosensor surface (‘AndromeDA’). **g**, Stimulation of the neuron leads to the release of DA, which interacts with AndromeDA. This interaction triggers the nanosensor fluorescence, allowing for the detection of the spatiotemporal pattern of DA release and diffusion (left), and to analyze release kinetics (right).

Offering a general solution for analyzing imaging-based exocytosis assays in a variety of cell types, acquired at diverse speed and with varying video signal quality has proven to be of great challenge. The largest source of variability comes from the cell type, the vesicle type and the method used to visualize exocytosis (**Fig. 1**). In general, exocytotic events can be classified into three main categories. The first two categories pertain to the measurement of vesicle exocytosis (**Fig. 1a-e**), viewed as a transient change of fluorescence intensity on discrete spots. Therefore, we call these events, fluorescent burst events (FBE) which can occur at either fixed or non-fixed positions. When measuring the exocytosis of individual vesicles, the vesicles often move towards the fusion site before exocytosis occurs (**Fig. 1a-c**). This site can be anywhere on the plasma membrane, making multiple fusions at the same site unlikely. These events are termed non-fixed FBEs. In contrast, synaptic transmission in neurons occurs exclusively at synapses and encompasses the fusion of multiple vesicles within a brief period, resulting in a change in fluorescence intensity at static positions. This is observed during spontaneous exocytosis as well as during repetitive stimulation (**Fig. 1d-e**). Thus, the analysis can be conducted without considering moving fluorescent objects. These events are referred to as fixed FBEs. Finally, a third category of events arise from secreted chemicals that are measured with nanosensor technology. In such events, the secreted chemical induces fluorescence changes across a relatively large area in the extracellular space in a concentration-dependent manner (**Fig. 1 f-g**). We call these events hotspot area events. The detection of all these different types of exocytic event require different analytical approaches.

We have developed an adaptive automatic vesicle fusion detection program, termed IVEA, pronounced [’Λɪvi], which uses deep learning to analyze a wide array of vesicle fusion events. It is a trainable modular structured tool that uses a hybrid approach based on the combination of computer vision and AI. IVEA introduces three new detection and recognition techniques for analyzing exocytosis events. The first two techniques involve discriminator neural networks to detect the exocytosis of fluorescently labeled vesicles in a spectrum of experimental paradigms. The first one detects dynamic and heterogenic fusion pattern of large vesicles in cells like T cells, INS-1 cells and chromaffin cells—in other words, non-fixed FBEs from any cell system. It is based on a vision transformer network (ViT) (**Fig. 1c**), which was chosen for its adaptability^21^. This neural network was trained to detect pH-sensitive (**Fig. 1e**) and pH-insensitive (**Fig. 1d**) labeled vesicle cargo secretion. The second has been developed for fixed FBEs detection and analysis (**Fig. 1a,b**). It is based on long short-term memory (LSTM) network^22^ due to its comparatively low computational resource demands (**Fig. 1a,b**). The third technique, which involves machine learning, is used for detecting and analyzing hotspot areas like those generated by nanosensor-based technologies such as AndromeDA^23, 24^ and composite nanofilm^25^ (**Fig. 1f-g**).

The event detection method employed by IVEA makes use of the grayscale image foreground detection method, which employs forward and backward subtraction. This facilitates the determination of the fluorescence intensity fluctuation associated with exocytosis within a specified time step. This back and forth subtraction enables the determination of the positive variations in the decrease and increase of fluorescence intensity, respectively. In contrast, in the absence of any fluctuations or foreground variation, image subtraction yields an image characterized by pixel values of zero. However, the presence of noise and/or artifactual fluctuations during image acquisition makes it difficult to differentiate real events from artifacts. To analyze the subtracted images, we select for each image regions of interest (ROIs) that are centered on local maxima (LM), which are then subjected to further classification. In the case of non-fixed FBEs, sequence of image patches is extracted, encompassing frames both preceding and following the time point of the event. Each sequence is fed into a ViT network, which classifies them as exocytosis or not. We have developed an encoder for our ViT architecture^21^, designated as eViT. The encoder network processes image patches of 32×32 pixels. Each patch represents the extracted area centered at the LM, while the sequence is centered on the fluorescence intensity peak time (**Fig. 2a**). The encoder network automatically extracts features from each sequence and forwards the encoded data to a multi layered perceptron (MLP), which in turn forwards the data to the ViT network. The ViT network then performs positional encoding on the extracted features and classifies each batch as a true event or not (**Fig. 2b**). For fixed FBEs, we employed a straightforward model architecture comprising a LSTM network^22^ for exocytosis classification. Our LSTM architecture is designed for multivariate time series classification^26^ (**Fig. 2c**). In this case, the image patches undergo preprocessing to extract features and convert them into time series data of one-dimensional time series vectors (**Fig. 2a**). The preprocessing stage involves the extraction of features from the sequence of image patches, with each patch divided into 13 regions. Each feature vector represents the average gray value over time for a specific region. These feature vectors are subsequently fed into the LSTM network for classification (**Fig. 2d**).

**Figure 2.**
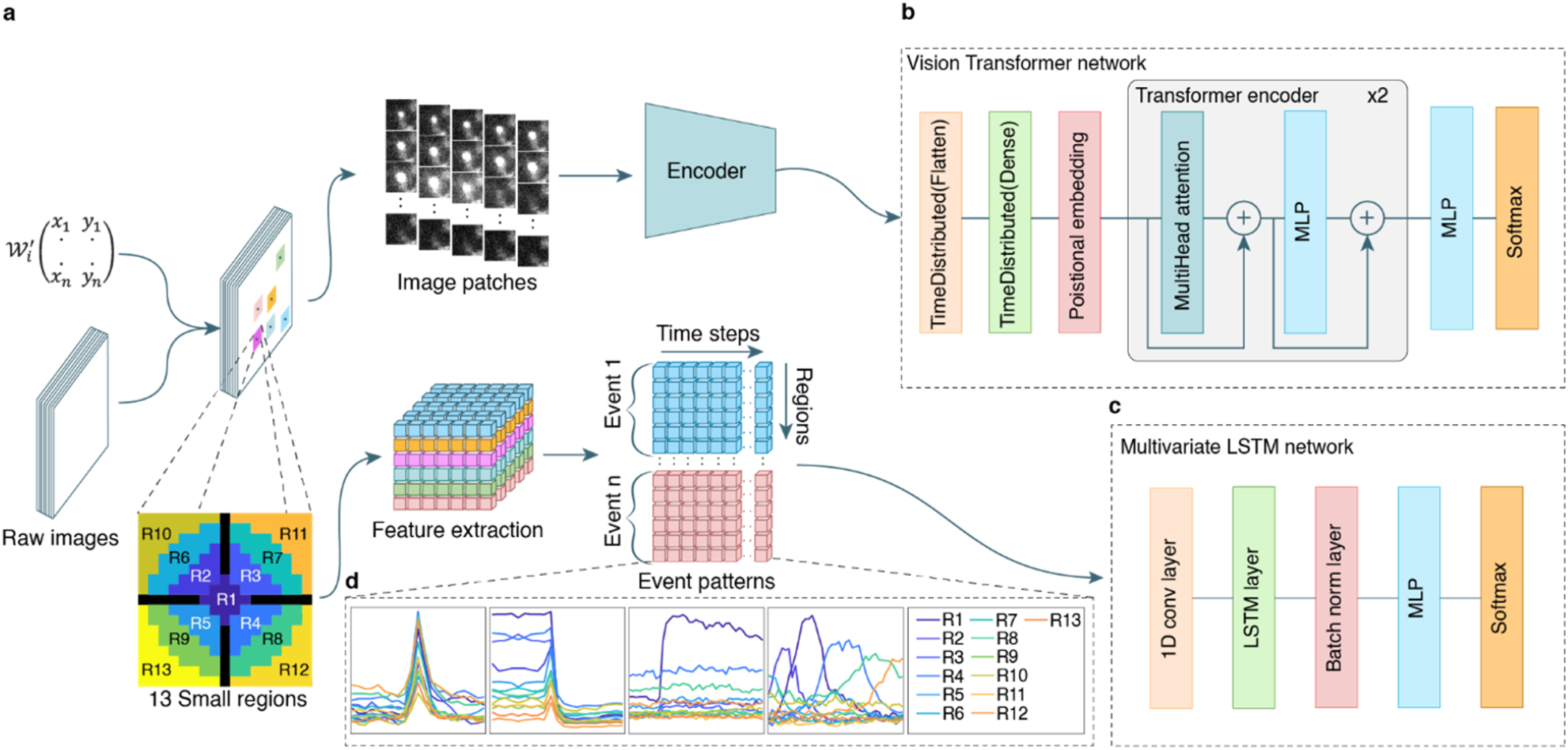
Overview, our neural networks and the feature extraction process. **a**, Flow of the two distinct methodologies employed to prepare the data prior to classification. LM coordinate vector 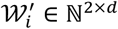 represent the selected region. In the case of non-fixed FBEs, the regions are extracted and then input to an encoder network. A total of 26 patches were extracted for each selected region of size 32×32 pixels. For fixed FBEs, feature extraction is performed. Feature vectors are represented as 13 regions, which are centered around the event’s local maxima (*R*_l_, *R*_2_ … *R*_l3_). The black regions in the “13 Small regions” scheme represent buffer zones. The features extraction for fixed FBEs are sorted as number of events, regions and times series. **b**, Vision transformer network architecture. **c**, Multivariate LSTM network architecture. **d**, LSTM network recognizes each package of data as a graph of 13 curves representing the regions normalized mean intensity variation over time. The LSTM model is available for non-fixed FBE analysis as well. Event patterns are representative graphs of 1. single T cell vesicle fusion in which the fluorescent cargo was either pH-sensitive (left) or 2. pH-insensitive (middle left); 3. fusion at neuron synapse in which the vesicles are stained with pH-sensitive membrane protein (middle right); 4. fusion of a single vesicle that moved (right).

IVEA possesses the capacity for automatic parameters estimation, a feature that facilitates and accelerates its utilization. The extraction of all image’s LM coordinates results in the selection of a huge number of LM coordinates including those at noise level. This will result in significant computational demands, particularly when utilizing the eViT network, due to the vast number of sequences of image patches that must be analyzed. The automatic parameters reduce the number of LM by estimating a sensitivity threshold and the average ROIs full width at half-maximum (FWHM) (see Method section).

In biological sciences, the utilization of machine learning, particularly deep learning, is experiencing rapid growth due to its reliability and effectiveness. It is applied in different areas such as image segmentation, data analysis or 3D model prediction^27-29^. Unlike mathematical models which can only detect specific events, the deep learning used in IVEA can learn and distinguish between multiple types of events. Additionally, IVEA can adapt to a diverse range of data, allowing for batch data analysis without human input. Our software is capable of collecting a broad amount of information, which allows robust statistical analysis that is superior to existing tools. To ensure IVEA’s wide applicability, we chose to run it through the broadly used and open-source Fiji platform (**Supplementary Fig. 1**). While IVEA is fully automated, users are offered the option to fine-tune the plugin’s advanced settings (**Supplementary Fig. 2**) or train the models to accommodate specific experimental paradigms.

## Results

Detection and accurate prediction are two critical factors for analyzing fusion events. To evaluate our plugin, we compared the IVEA detection and prediction with human experts (HE) manual detection. Our research was conducted using diverse datasets, from multiple laboratories and different microscopes, each belonging to one of three main categories. For the first category, we analyzed the exocytosis of lytic granules in T cells and of dense core vesicles in INS-1 cells and in chromaffin cells. To broaden the range of applications, we used cells in which the granules were stained either with a pH-sensitive or pH-insensitive fluorescent protein. The second category was centered on the examination of dorsal root ganglia (DRG) neurons expressing Synaptophysin-SEP (SypHy), which enabled the investigation of synaptic transmission at neuronal synapses. Lastly, our software was used with dopaminergic neurons surrounded by a surface ‘painted’ with dopamine nanosensors (‘AndromeDA’), which allows the study of dopamine release events. Most of the data was analyzed with IVEA using the default settings and automated parameters. We used videos that were devoid of events within the first second of acquisition as these frames are used by IVEA to learn these automated detection parameters. However, users can opt out of automated learning in favor of manual override. The IVEA analysis was conducted on a range of computer systems, with a baseline configuration of an Intel Core i5 processor and 32 GB of RAM without GPU.

### Event simulation validation

We evaluated our software performance by creating simulated videos with a ground truth that emulates the fusion of lytic granules in CTL. Subsequently, IVEA batch analysis was conducted on the aforementioned videos using average computers without relying on GPU (see Methods). We employed IVEA’s default parameters with the exception of the neural network radius, which was set to 16 pixels. All simulated fusion occurrences were successfully identified without any false positive detection. To ascertain the limitations of IVEA accuracy, we have introduced white noise and Poisson over the videos (**Fig. 3a**).

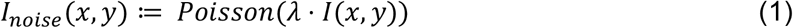

The Poisson noise scaling factor λ was increased from 0.1 up to 10 times the signal. We run IVEA on our simulated noise enriched videos and achieved recall of 99.71% ± 0.29%, a precision 94.49% ± 3.23% and F1 score of 96.71% ± 1.91% for λ = 0.1 up to λ = 1, which corresponds to a rather low SNR (**Fig. 3c**). When λ was increased further, IVEA began to lose some of the small vesicle, due to the added Poisson noise that surpassed the signal strength. For λ = 1 up to λ = 10, the recall was 96.86% ± 2.55%, a precision was 79.22% ± 4.68% and F1 score was 86.51% ± 3.27%. However, these are noise levels exceeding experimental TIRFM videos, which would be analyzed by the scientist. Our results show that IVEA is ready to analyze real data, acquired with a challenging signal-to-noise ratio.

**Figure 3.**
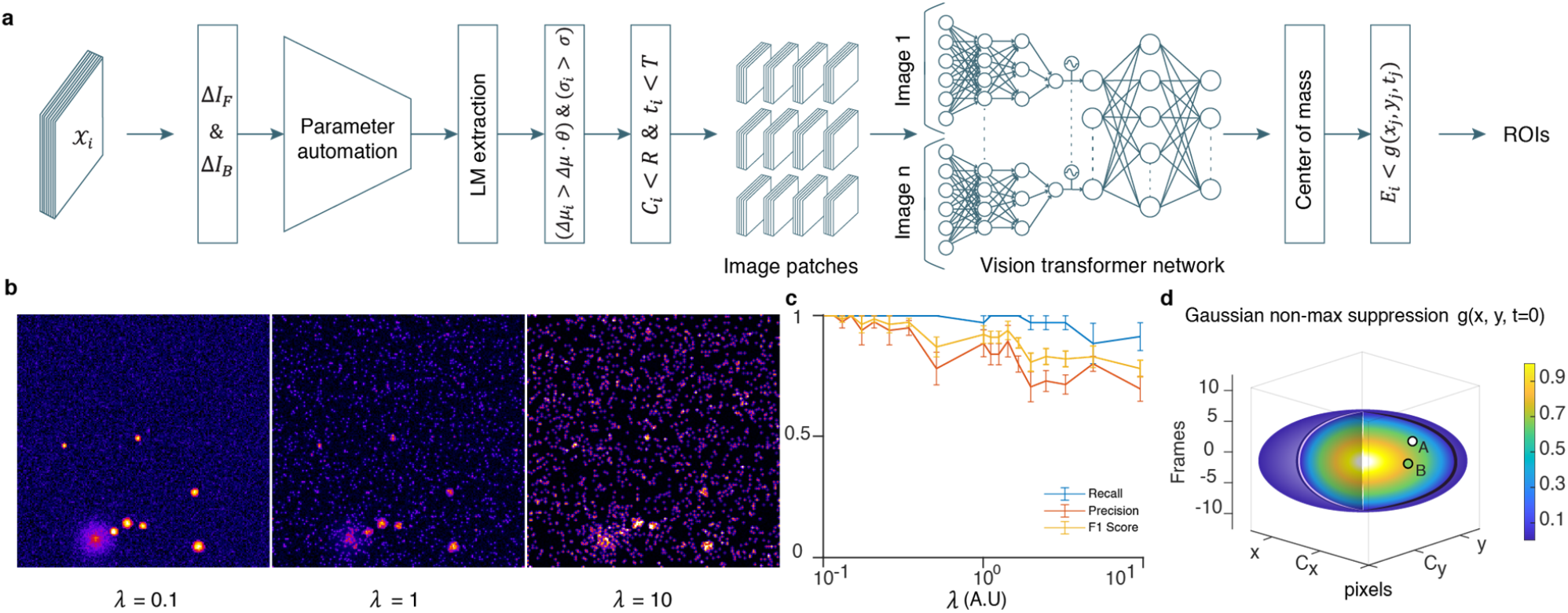
Overview of non-fixed FBE detection algorithm and simulation analysis. **a**, Algorithm flowchart, 𝒳_*i*_ is the image sequence. Δ*I*_*F*_ & Δ*I*_*B*_ are the forward and backward subtraction images Δ*I*. Δ*μ*_*i*_, *σ*_*i*_, *C*_*i*_ and *t*_*i*_ are the event *E*_*i*_ mean intensity, FWHM, center coordinate and the time for which *E*_*i*_ was detected, respectively. Δ*μ* and *σ* are the automated parameter for detection. *θ, R* and *T* are the detection sensitivity, search radius of 3 pixels and the time interval of 4 frames, respectively. Image patches are the 32×32 pixels extracted over time for each selected region. The network scheme is an encoder-vision transformer network. The “Center of mass” step is a process of adjusting the center of mass of true positive events. The final step of the algorithm is the Gaussian spatiotemporal function, which is used as a non-max suppression method. **b**, Effect of Poisson noise added to simulated videos. Left: simulated video with Poisson noise scaling factor λ = 0.1 with an ideal exocytosis event in a cytotoxic T-lymphocyte. Middle: same video with λ = 1. Right: same video with λ equal 10 times the signal. **c**, The graph presents the evaluation of our eViT performance following the analysis of simulated videos with a noise scaling factor that varied between 0.1 and 10. The graph shows the average recall, precision, and F1 score values with SEM (n=5). **d**, 3D ellipsoid representing the Gaussian spatiotemporal search equation *g*(*x*_*j*_, *y*_*j*_, *t*_*j*_) for event *E*_*j*_. The color bar ranging from 0 to 1 corresponds to the pixel mean gray value at point (*x, y, t*). Point A and B are two true positive events occurring very close in time and space to *E*_*j*_. Event A mean gray value at point (*x*_*A*_, *y*_*A*_, *t*_*A*_) is greater than *g*(*x*_*j*_, *y*_*j*_, *t*_*j*_) which means A is a separate event. While event B mean gray value at point (*x*_*B*_, *y*_*B*_, *t*_*B*_) is less than *g*(*x*_*j*_, *y*_*j*_, *t*_*j*_), which means *E*_*j*_ =B.

### Non-fixed FBEs analysis

We have analyzed several videos in which CTL secreted lytic granules (**Fig. 4 a-b** and **Supplementary Video 1-4**). In CTLs, lytic granules were stained via expression of granzyme B, a cargo protein, that was tagged with a fluorescent protein. This fluorescent protein was either the pH sensitive pHuji (**Fig. 4a**) the weakly sensitive eGFP or the pH insensitive tdTomato (**Fig. 4b**) generating very distinct event signatures as displayed in Fig. 1d-e. The videos were recorded at 10 Hz over 10 min and encompassed 1 to 30 CTLs. In these videos the total number of fusion events detected by the HE was around 234 (**Fig. 4d, Supplementary Video 1-4**). The time that the HE required to analyze one cell was 10 min for lytic granules labelled with pH-insensitive fluorescent protein and 5 min for lytic granules labeled with a pH-sensitive fluorescent protein. This amounts to 300 minutes of analysis for videos containing 30 cells. IVEA required less than a minute per cell with a video size of ∼256×256 pixels and 300 frames. On a computer equipped with an Intel Core i9 10^th^ generation, this sums up to about 15 min for the entire video with 30 cells irrespective of the fluorescent marker protein. The performance of IVEA platform was further evaluated on a CTL dataset acquired in a separate laboratory equipped with a different TIRFM. The lytic granules were stained using LysoTracker Red thereby displaying different fusion kinetic (**Supplementary Fig. 3c**). To increase signal diversity, we also analyzed non-fixed FBEs in chromaffin cells (**Fig. 4c**) and INS-1 cells (**Fig. 2d**) that secrete dense core vesicles. In these cells the vesicles are smaller than in CTLs and they are often more packed. The vesicles were stained via the over-expression of NPY attached to a fluorescent marker that are weakly pH sensitive like mGFP or mNeonGreen or pH insensitive such as mCherry.

**Figure 4.**
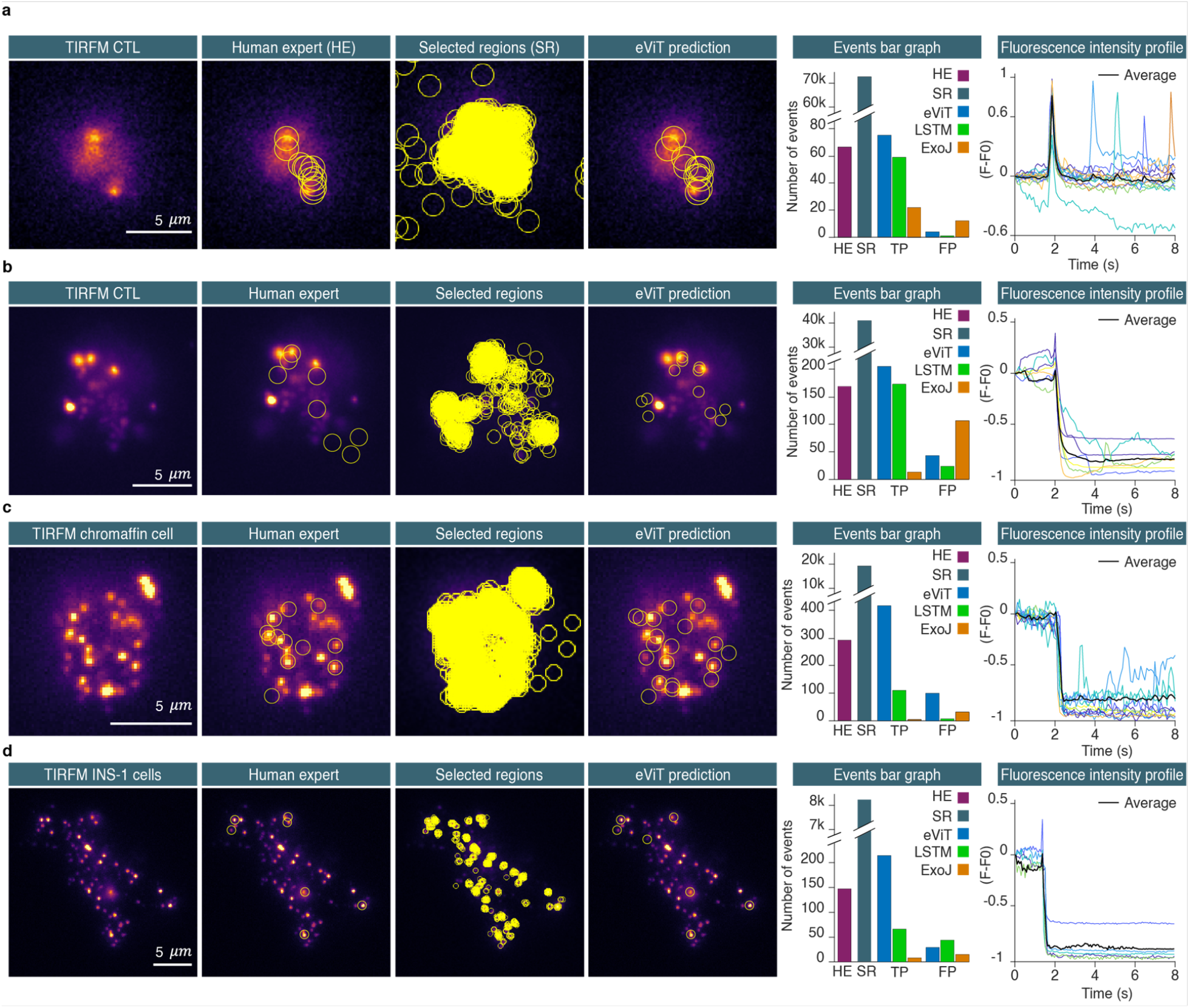
Non-fixed FBE analysis. Detection process compared to human expert (HE) displayed from left to right, ROIs are shown as overlay. **a – d**, from left to right, the first panel shows the TIRFM raw image of a cytotoxic T cell, chromaffin cells and INS-1 cells. The second panel shows ROIs of the detected events by the HE. The third panel represents the raw image with all the selected regions prior to classification. The fourth panel shows the classified events by our ViT network. The bar graphs show the total number of events identified by the HE (purple), the number of selected regions (SR, gray), the number of classified events by the ViT (blue), the LSTM (green), and ExoJ (orange). Shown are the average results of all analyzed TIRFM videos. The line plot panel shows the event fluorescence intensity profiles of the true positive events detected by IVEA. The events profiles are aligned on their respective detection time. The fluorescence peaks at time point 2 sec corresponds to the exocytosis of the detected events. **a**, CTL expressing pH-sensitive granzyme B-pHuji. 10 movies of individual cells were analyzed (n_cell_=10). **b**, CTL expressing pH-insensitive granzyme B-tdTomato. 7 videos containing 1 to 11 cells were analyzed (n_cell_=33). **c**, Chromaffin cell expressing NPY-mCherry (pH insensitive fluorescent protein). 5 videos were analyzed which were pooled together with 5 videos of INS-1 cells expressing NPY-mCherry (n_cell_=10). **d**, INS-1 cells expressing NPY-mNeonGreen. For the analysis, 9 videos of individual INS-1 cells expressing NPY tagged to mGFP or mNeonGreen were pooled together as all fluorophores were weakly pH sensitive but showed distinct cloud release (n_cell_=9). INS-1 cells videos were acquired at the Medical Cell Biology, Uppsala University, Sweden.

The eViT network for non-fixed FBEs was trained with 10 distinct categories. The event categories encompass: two distinct types of exocytosis events, fusion with a cloud and fusion without a cloud (sudden disappear); and 8 other types of events, such as fast drift or focus change, vesicle movement, random noise, random noise with intensity fluctuation, vesicles with noise, and vesicle docking and undocking. Analyzing the videos by HE identified 673 fusion events in all videos, across all cells and labels. IVEA detection routine registered around 142k ROIs for later classification, out of which 2239 were identified as true events by our eViT network. Non-fixed FBE can exhibit spatial spreading of fluorescence intensity (**Fig. 1d-e, Supplementary Video 1-4**), which leads to multiple detection of the same events (duplicates). Various non-maximum suppression techniques are typically used to address this problem, including the classical intersection over union (IoU) method, weighted boxes fusion (WBF), and others^30^. However, these methods cannot be implemented with our data because exocytotic events cannot be limited to objects with boundaries or boxes. Thus, we have developed a new method that implements 3D Gaussian spread over time (**Fig. 3d, Eq. 2**).

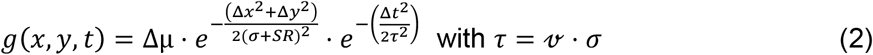

where Δ*μ* is the fluorescence intensity of the event’s area over subtracted image, *σ* the event’s cloud spread, *SR* = 1 the spread radius controlled by the user, and event temporal cloud spread *τ* and 𝓋 the image acquisition frequency set to 10 Hz.

To find prime events from the redundant ones, we apply these criteria using *g*(*x, y, t*) on both events such as:

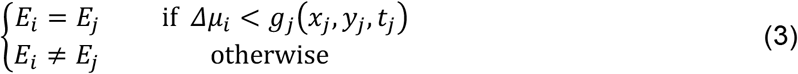

where *E*_*i*_ and *E*_*j*_ are two different true positive events.

After applying our non-max suppression algorithm, 919 true positive events were selected while 1,320 duplicates were discarded. Therefore, IVEA detected 246 additional events from those identified by the HE.

We compared our two neural networks, the eViT and the LSTM which were trained on the non-fixed FBEs for detecting exocytosis (**Fig. 2a**). All true positive events identified by the eViT, and the LSTM were verified by the HE. The analysis was conducted on the same sets of videos as previously described. The results were divided according to the cell type and vesicle label.

The first set represents CTL with pH-sensitive staining (**Fig. 4a**). The results yielded a total of 74 true positive events identified by the eViT and 59 by the LSTM, in comparison to 66 events identified by HE. The eViT achieved a recall of 100.00% ± 0.00%, a precision of 96.73% ± 1.60% and an F1 score of 98.27% ± 0.85%. The LSTM was performing similarly to the eViT with a recall of 84.29% ± 5.38%, a precision of 97.62% ± 1.65% and an F1 score of 89.57% ± 3.67% (see supplementary Tabel 1 for statistics).

For set two which represent CTL with pH-insensitive staining (**Fig. 4b**), the eViT detected 219 true positive events, while the LSTM identified 172, in comparison to 168 events found by a HE. The eViT achieved a recall of 100.00% ± 0.00%, a precision of 81.06% ± 3.92% and an F1 score of 89.18% ± 2.35%. The LSTM demonstrated a sight but not significant reduction of the recall value of 89.14% ± 4.78%, a precision of 88.89% ± 2.32% and an F1 score of 88.48% ± 2.79% (see supplementary Tabel 1 for statistics). The eViT also detected proficiently exocytosis of lytic granules stained with LysoTracker Red (**Supplementary Fig. 3**) with an average recall of 100% ± 0%, a precision of 66.59 ± 6.24 and a F1 score of 79.27 ± 4.50 (n_cell_=4).

The third set is for smaller cells with challenging exocytosis features, chromaffin cells and INS-1 cells stained with pH-insensitive probes (**Fig. 4c, Supplementary Fig. 4**) in which the eViT performed significantly better than the LSTM (see supplementary Tabel 1 for statistics). The eViT detected 412 true positive events, the LSTM detected 110 compared with 292 events identified by a HE. The eViT demonstrated a recall of 98.21%± 6.64%, a precision of 78.27%± 3.48% and an F1 score of 86.58% ± 9.81%. As for the LSTM showed a recall of 27.74% ± 6.64%, a precision of 93.49% ± 3.48% and an F1 score of 41.67% ± 2.79%.

For the last set which represent INS-1 cells stained using NPY-mNeonGreen and NPY-mGFP, the eViT identified 214 true positive events, in comparison to 66 by the LSTM, while the HE detected 147 events. The eViT recorded a recall of 98.70%± 0.93%, a precision of 90.08%± 2.54% and an F1 score of 94.05%± 1.53%. In contrast, the LSTM showed significantly reduced recall of 29.89% ± 4.35%, a precision of 78.50% ± 8.67% and an F1 score of 40.08% ± 4.63% (**Fig. 4d**, see **Supplementary Tabel 1** for statistics).

Overall, the average results across all sets show that the eViT outperforms the LSTM network, as the eViT achieved an average recall of 99.23% ± 0.40%, with average precision of 86.53% ± 3.66% and average F1 score of 92.02%± 2.25%. In contrast, the LSTM had an average recall of 57.76% ± 14.50% and average precision of 89.62% ± 3.56% and average F1 score of 64.95% ± 12.04%. Therefore, we chose the eViT as the main model for the IVEA software for the non-fixed FBEs analysis while the LSTM is still can be chosen by the user.

IVEA was compared to an existing ImageJ open source plugin called ExoJ^18^. While ExoJ is designed to detect pH-sensitive FBEs, we subjected ExoJ to analyze both pH-sensitive and insensitive staining and compared the results with those obtained using eViT. As expected ExoJ didn’t work properly when the vesicles were labeled with pH-insensitive probes (**Fig. 4b, c, d**). For a pH-sensitive probe (**Fig. 4a**), ExoJ detected 22 events using the program’s default settings, showing a recall of 29.52% ± 6.15% and a precision of 71.63% ± 9.88% with an F1 score of 39.41% ± 6.51%, in our hands. These values are significantly lower than those obtained by the eViT (see **Supplementary Tabel 1** for statistics). These 22 events exhibited a distinctive fluorescence intensity profile, which corresponded to constraints imposed by the algorithms that were implemented in ExoJ^18^. Iterative adjustments of ExoJ parameters improved the number of detected events at the cost of false positive events (**Supplementary Table 2**). Furthermore, this iterative adjustment has to be performed of each individual movie due to diverse nature of the fusion events present in our data. Additionally, we tested pHusion another program based on mathematical model^31^. This program did not yield better results than ExoJ (see **Supplementary Table 3**). In contrast, our deep learning-based technique exhibited a superior performance, effectively capturing the variety and complexity of the fusion events that exceeded the scope of certain mathematical model algorithms.

IVEA output consists of two files sets: an ImageJ ROI zip file and an analysis CSV file. Each ROI is labelled and positioned on the center of mass at the peak of the fluorescence intensity of the event to which it corresponds. The CSV file contains measurements of the fluorescent intensity of each event over a fixed time interval.

### Fixed FBEs analysis

Fixed FBEs were analyzed using the second branch of our platform, which employs the LSTM neural network for classification (**Fig. 2a, c**). Our eViT for non-fixed FBEs requires an input sequence of 26 image patches with a size of 32×32 pixels. This necessitates substantial memory and computational power both during the training phase and classification task. In the case of fixed FBEs, the number of frames per sequence required to analyze an exocytosis event needs to be significantly higher than for non-fixed events due to different signal kinetics. A reasonable number of frames to study them would be between 41 to 100 frames. Furthermore, the number of selected regions to extract the image patches is high, with up to 14k ROIs in a single video. This would result in huge memory demands, necessitating the use of high-end computational resources with our current eViT to analyze fixed FBEs.

We analyzed DRG neurons videos expressing SypHy that were derived from Shaib *et al*.^32^ and Staudt *et al*.^33^. These videos display a variety of synapse count, intensity, vesicle movement, and background activity. In our data set, fixed FBE were characterized by fast rise (within 4.1 s, i.e. 41 frames acquired at 10 Hz, **Supplementary Fig. 5d**) of the fluorescence intensity in a spot like area (**Supplementary Fig. 6a**). Our neural network input layer for the fixed FBEs was adapted for the input vector 𝒫(*X*(*t*), 𝔣) ∈ ℝ^𝔣×T^, where *X*(*t*) ∈ ℝ^T^ as 𝔣 is the number of regions and *t* is the time series for 0 < *t* ≤ *T* = 41 with *t* ∈ ℕ. Thus, the LSTM network discards the majority of 𝒲^′^ (i.e. the selected regions, **Fig. 2a**) resulting in the classification of highly probable true events. For events in which the rise time was longer than 41 frames, the LSTM network sorted the events into the “intensity rise” category (**Supplementary Fig. 5c**) thereby discarding them from the collection of true events. For experimental conditions in which long lasting events with slow kinetics are the result of long stimulation paradigms or very high acquisition frequency (**Supplementary Fig. 6**), we adapted an “add frame” option that allows the user to adjust the event’s time window by increasing the number of frames by *t*_*n*_, such as *X*′(*t*) ∈ ℝ^𝓉^ with 𝓉 = *T* + *t*_*n*_. To determine 𝒫(*X*(*t*), 𝔣) ∈ ℝ^𝔣xT^, the extended measurement is then sampled using windowing mean sampling method such as:

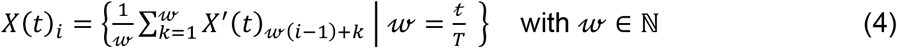

Where, *X*(*t*)_*i*_ is the *i*-*th* element of the sampled vector *X*(*t*), and *X*′(*t*)_*j*_ as *j* = 𝓌(*i* ™ 1) + *k* is the *j*-*th* element of vector *X*^′^ (*t*).

The videos that were analyzed, were acquired at 10 Hz and comprised 3000 frames, each measuring either 512 by 512 pixels^32^ or 512 by 256 pixel^33^. The DRG neurons were stimulated electrically for 1 min^32^ or they were stimulated twice for 30 s and 1 min with 10 s recovery phase in between^33^. Due to the different stimulation paradigms, the second video datasets yielded more long-lasting events. The human analysis of these videos was a challenging task that required an average of 60 minutes per video. Detected events had predominantly a high fluorescence intensity variation or lasted for relatively long periods of time. IVEA reduces the time of analysis of the same videos to under one minute per video. Furthermore, batch analysis capabilities exclude the need for manual parameter adjustment, as the tool autonomously learned and adapted to the input video’s characteristics. The results of the analyzed videos show that the neural network was able to classify virtually all the human labeled regions (**Fig. 5b,c**). Additionally, the neural network was able to detect more true fusion events than HE had originally detected by double checking and validating them as real events. The HE was able to detect overall 356 fusion events, while IVEA detection routine registered 84k ROIs for later classification. Most of these events were identified by the neural network as false events. A total of 2049 events were classified as true events, while only 70 events were identified as false positives (**Fig. 5c**). The average recall, precision and F1 score were 88.12% ± 2.70%, 96.37 ± 0.45%, and 91.83 ± 1.61% respectively. Importantly, in comparison to the HE approaches, the LSTM network for fixed FBEs could detect weak events or events with fast kinetics (**Supplementary Fig. 5e,f**; **Supplementary Video 5**). Conversely, the HE was able to detect events with very slow kinetics (**Fig. 5b**) (**Supplementary Fig. 6b**). Due to the “intensity rise” category (**Supplementary Fig. 5c**) these slow evens (i.e. longer than 41 frames) were missed by IVEA in the videos from Staudt *et al*.^33^. However, they were detected when applying the “add frame” option that extends the time interval by adding 60 frames for correct classification (**Supplementary Fig. 5, Supplementary Video 5**). Overall, IVEA identified about 5.4 times as many true events as HE. We compared IVEA to the existing open source software SynActJ^20^ that was devised to analyze synaptic activity in neurons stained with the overexpression of synaptobrevin-SEP or alike proteins. First, we compared the result of IVEA and SynActJ on the provided test movie. SynActJ and IVEA were able to identify most of the active synapses but missed one and yielded one false positive event (**Supplementary Fig. 7a**). Then, we compared both programs on one of our movies in which synaptic activity was clearly detectable by HE. While IVEA detected all events without any false positive events, SynActJ was able to detect only a very limited number of active synapses and showed a significant number of false positive events (**Supplementary Fig. 7b**). Thus, IVEA is by far superior to SynActJ.

**Figure 5.**
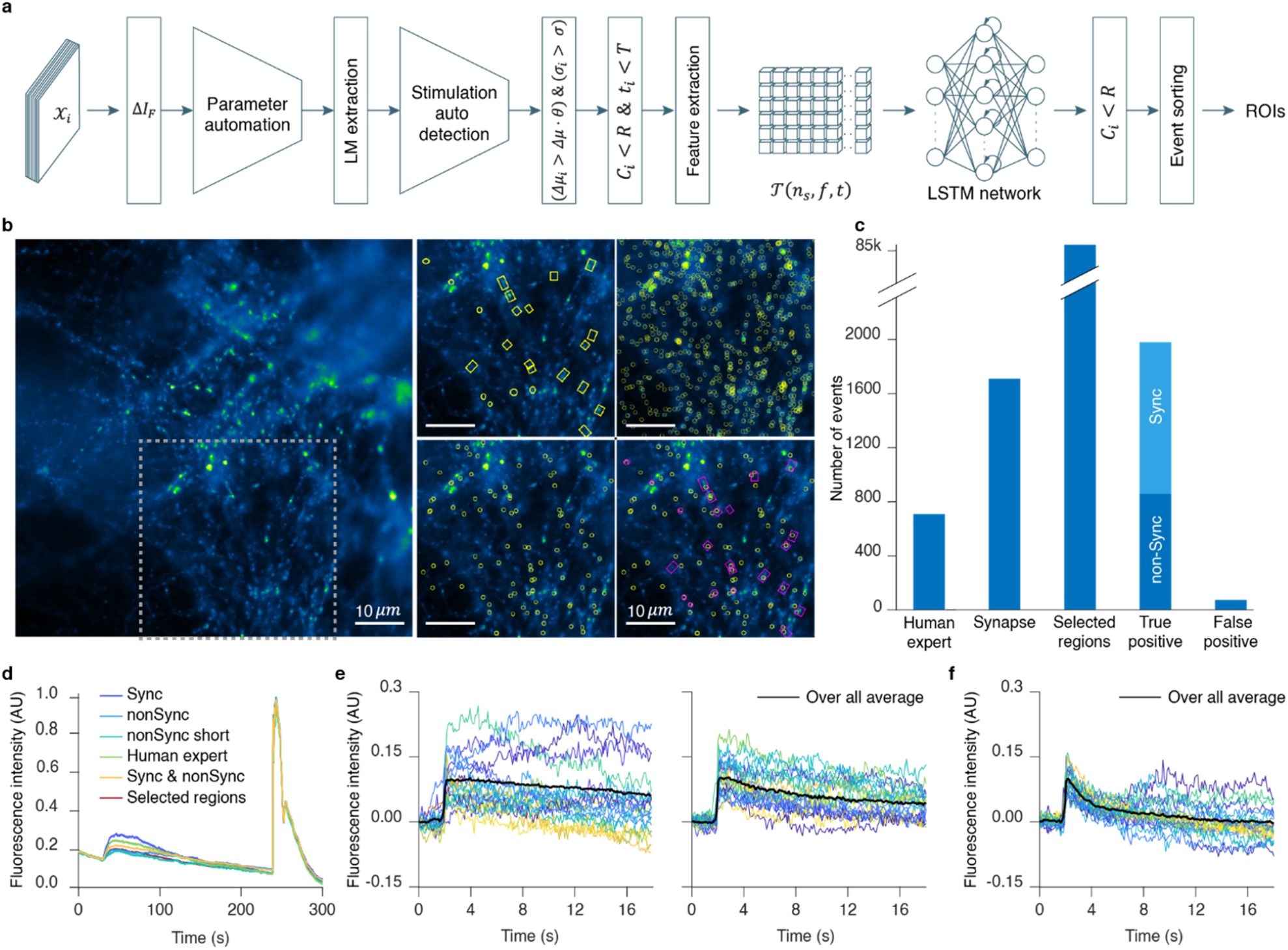
Detection and analysis of exocytosis in neurons with fixed FBE Algorithm. **a**, This panel displays the fixed FBEs algorithm flowchart. Defining parameters: 𝒳_*i*_ is the raw images; Δ*I*_*F*_ is the forward subtracted image. Δ*μ*_*i*_, *σ*_*i*_, *C*_*i*_ and *t*_*i*_ are the event *E*_*i*_ mean gray value, FWHM, center coordinate and the time of the event; Δ*μ* and *σ* are the threshold mean gray value and FWHM. *θ, R* and *T* are the detection sensitivity, search radius and the event’s time interval. 𝒯(*n*_s_, 𝔣, *t*) is the extracted data in 3D tensor shape. **b**, Left, raw image of DRG neurons over-expressing SypHy forming synapses on spinal cord neurons. Right, depicts the area within the dashed gray box on the raw image overlayed with 4 different ROIs. Displayed are from left to right, top to bottom, the human expert (HE) ROIs, the selected regions (SR) ROIs, the neural network ROIs, and a comparison of HE (Magenta) and neural network (Yellow) ROIs, respectively. **c**, Bar graph representing the total number of events analyzed in 11 DRG neurons videos. IVEA parameters for the analysis were set to default. **d**, Overall mean intensity profile of the combined ROIs areas comparing different events types. **e**, Mean intensity profile over time representing the events detected at the stimulation time (synchronized events), the second graph is for the events detected before or after stimulation (non-synchronized). **f**, Mean intensity profile for short events category whether it was synchronized or non-synchronized. The event intensity profiles are aligned on their respective detection time (**e & f**).

For advanced analysis, IVEA distinguishes between various event types and categorized them based on their timing in respect of the experimental stimulation paradigms. The events can be classified as synchronized to the stimulus or unsynchronized (**Fig. 5c**). This feature was implemented as both types of events might show differences in kinetics. Stimulation time can be set manually, but to ease usability we also implemented an automatic stimulation detection. Our neural network was trained on *n*_*c*_ = 9 distinct events categories. These categories include: four types of events (fusion, short time fusion ∼4 frames (0.4 sec at 10 Hz), electrical or agonist stimulation, and NH_4_^+^ stimulation) and five types of artifacts (fluorescence intensity rising, out-of-focus artifacts, vesicle motion, white noise, and fluorescence intensity fluctuations).

While events are sorted and labeled, spatial recognition task is performed to locate and unify the event’s spatial identity while counting how many times the same synapse was active. Finally, the output for the fixed FBEs are ROIs files whereby each ROIs is positioned at the event’s maximum intensity temporal occurrence. The ROIs are fully labeled with an ID, frame number and event status. Additional outputs are the summary for each event and the measurements, such as intensity over a specified time interval and over the full video length.

### Hotspot area extraction

For the third analysis conducted using IVEA, we employed distinct techniques from those mentioned earlier. In this analysis, we assumed that the sensor array was in a fixed position, awaiting the occurrence of hotspots. Due to the limited availability of training data and the simpler features as fixed and non-fixed FBEs, we opted not to implement a neural network. Instead, we utilized machine learning and iterative thresholding for detection, and employed spatial search and mean intensity tracking over time for event recognition (**Supplementary Fig. 8a**). Before applying the foreground detection method, we perform intensity fluctuation correction. The challenge while correcting the intensity fluctuation is to avoid altering and distorting the event as much as possible. To address this, we have developed a new method, which we call “Multilayer Intensity Correction” (MIC) (**Supplementary Fig. 9**). The main idea of our method is to perform pixel value correction based on the variation of the average value of each cluster of pixels. To preserve the signal intensity, the signal should be a small range of pixels registered in a cluster, otherwise the signal is affected by the average value adjustment. MIC algorithm is performed through segmenting a given image into *k* layers using the k-mean clustering algorithm^34^ after performing Gaussian filter of sigma equal to 1. After performing foreground detection, we extracted the hotspots from the processed image Δ*I* by converting it to a binary image using a global threshold (**Supplementary Fig. 8b**). Since Δ*I* is the resulting matrix of intensity variation between two images, applying different types of threshold algorithms specially those found in Fiji, lead to unpredictable results. Therefore, we implemented our own iterative global threshold. The iterative threshold is not dependent on the statistical information calculated from the image. Instead, it is iterated over the noise level Δ*I*, with the objective of eliminating it (**Supplementary Fig. 8b**). This allows us to determine the global threshold of the fluorescence intensity value (see material and methods). After detection of the possible hotspot, the fluorescence intensity of each event is temporally tracked (**Supplementary Fig. 8d**). When the fluorescence intensity of the event falls below the mid intensity, the event signal is considered to have disappeared and the tracking stops (**Supplementary Fig. 8e**). The IVEA software, which uses advanced algorithms and automated parameters, reduced the need for users to iteratively adjust the parameters for analysis. Furthermore, it enhances the precision compared to the previous DART software using default parameters (**Supplementary Fig. 8f**).

## Discussion

IVEA is a robust pre-trained plugin for detecting and analyzing vesicle exocytosis. It seamlessly integrates into the Fiji platform for open-source accessibility. IVEA achieves an impressive performance with F1 scores as high as 98.27% ± 0.85% for the eViT (non-fixed FBEs, **Fig. 4**) and 94.46% ± 0.57% for the LSTM neural network (fixed FBEs, **Fig. 5**). Furthermore, IVEA is proficient in distinguishing real events from artefacts such as photon shot noise or fast transient focus change (**Supplementary Fig. 5a**). Three different algorithms can be selected by the user, making IVEA highly compatible with a range of exocytotic events displaying varying fluorescent intensity profiles. Additionally, IVEA’s high adaptability results from its machine learning foundation, particularly, leveraging deep learning ViT^21^, CNN and sequential models^35^. This enables IVEA to not only identify a wide range of exocytosis events, but also to learn and recognize new patterns, thereby expanding the scope of its capabilities.

Other Fiji-based exocytosis analysis plugins such as PTrackII^19^ SynAct^20^ or ExoJ^18^ have been developed. Their algorithms rely on fixed mathematical and/or morphological models that are proficient at detecting a limited type of events as these algorithms cannot adapt and learn by themselves. The detection parameters can be adapted to a certain degree by adjusting complex input parameters that can be overwhelming for user with limited programing skills. This also precludes easy implementation of batch analysis when the analyzed videos have different characteristics, such as varying noise levels or fusion kinetics. Other programs like pHusion, have been developed but are only available in MATLAB program, Python or other proprietary platforms making them less usable^15-17, 31^. Finally, an algorithm has been developed with the variability of the signal in mind, using CNN^14^, which is not made available as a software. In contrast, IVEA – a “plug and play” plugin, distinguishes itself by the adaptability of the detection and classification capability that is based on multivariate LSTM^26^ and ViT^21^ models. For fixed FBEs, we chose LSTM network for the analysis, as it requires less memory usage when compared to eViT. This is particularly important as the number of extracted sequence patches and the number of frames analyzed per sequence is high. An analysis of all these sequences by eViT would require huge computational resources that cannot be provided by CPU computation and a reasonable RAM size (about 64 GB). In contrast, for non-fixed FBEs, the presence of motion and other variables makes the classification of the events more complex. This necessitates the use of more sophisticated models, such as the ViT. To our best knowledge, this is the first time a software uses LSTM or ViT network for detecting and classifying a wide number of different fusion events.

To assess IVEA’s versatility, we extensively trained and evaluated it on exocytotic events in CTL, chromaffin cells, INS-1 cell and DRG neurons. Consequently, even with low SNR and presenting minimal features (dense core vesicles in INS-1 cells labeled with the pH-insensitive NPY-mCherry), we were able to detect exocytotic events with a high effectiveness, achieving a recall of 98.45% ± 3.79% and F1 score of 90.31% ± 5.67%. Additional challenges such as vesicle clustering do not impair IVEA’s capability to identify weak exocytotic events (**Supplementary Fig. 3**). Finally, IVEA successfully detected exocytotic events with different signatures than those used for training, albeit with some additional false positives (**Supplementary Fig. 4**). However, IVEA is not designed to detect non-burst events such as those recorded in hippocampal neurons labelled with Synaptobrevin-SEP. Indeed, in this case stimulated exocytosis at synapses result in long lasting increased fluorescence that spreads out of the synapse^36, 37^. While IVEA is virtually universal for detecting burst exocytosis in a wide array of experimental paradigms with the current trained models, users might still encounter specific needs. Therefore, we provide a Python app with a user-friendly GUI for training purposes (**Supplementary Fig. 10**). The choice of using Python was rooted in challenges related to vectorized computation, alongside variations in the versions of C++ jar files across Fiji, Google TensorFlow, and the deeplearning4J library.

IVEA shows robust analysis capability even when the noise power is equal to that of true events by extracting spatiotemporal features of the signal (**Fig 3c**). Additionally, IVEA is capable to discard artifacts such as short focus changes resulting in transient signal variation. However, slow focus drift cannot be compensated as well as lateral drift in the case of fixed FBEs. Upon drift an active synapse is detected at different locations and is assigned to several coordinates. Therefore, drift should be corrected using free plugins such as but not limited to NanoJ core^38^ prior to IVEA analysis. Non-fixed FBE does not require drift correction as the algorithm does not require fixed spatial coordinates. Finally, to detect exocytosis events, the IVEA algorithm requires to analyze the first 6 frames from each video. This step is important for the automation process, so that IVEA can learn from the images that are devoid of events (see method section “Fixed and non-fixed FBEs algorithm”). In the rare cases in which the videos contain events within these frames, IVEA can generate learning parameters that enable the detection of high SNR events while events with low SNR maybe overlooked. However, manually reducing the detection threshold to 1 or less can enhance the sensitivity of event selection. The neural network can classify these increased detections correctly, while eliminating most of the false positive events. As a result, a heightened computational time requirement may arise. This can be mitigated by extracting six frames that are devoid of events and adding them to the beginning of the movie.

## Conclusion

IVEA reduces analysis time by more than 90%, requiring minimal to no human input, and significantly decreases analysis bias. Furthermore, the number of fusion events detected by IVEA is higher than those detected by the human expert, especially when it comes to visualizing rare, i.e. low brightness, events in low SNR. This allows us to generate large datasets with meaningful statistical power. In addition, IVEA, unlike the human expert, readily detects short events, which not only increases the number of detected events but also promotes the understanding of biological processes. The earlier application of hotspot area extraction module for AndromeDA, revealed not only discrete dopamine (DA) release, extracellular DA diffusion, but it also enabled the discovery of heterogenous release hotspots - albeit using less advanced algorithm at the time. The trove of information collected from individual events allowed to elucidate the role of key proteins in exocytosis molecular machinery in dopaminergic neurons^24^.

We foresee that our program will be adaptable to many other exocytosis systems and beyond. For instance, the “hotspot area extraction” or the eViT method might be proficient at analyzing transient localized calcium signals with low SNR such as calcium sparks. Using IVEA for detecting exocytosis events has the potential to rapidly advance research in the neurosciences, immunology and endocrinology fields.

## Methods

### Fixed and non-fixed FBEs algorithm

Our approach of detecting FBEs involves two main parts: automatic region selection and neural network classification. The initial step is to determine the gray value variations in an image stack using foreground detection method. This is achieved by using a fixed sliding window that encompasses, by default, four frames in which the program performs forward subtraction and backward subtraction to form new 16-bit images representing the intensity variation Δ*I*_*F*_ and Δ*I*_*B*_. Backward subtraction was employed only with non-fixed FBEs to detect fusion events visualized with a pH-insensitive staining. After foreground detection method comes the detection parameters automation. This step is important to minimize user inputs and adapt the algorithm to better analyze each video. The program works under the assumption that in the first four images no events occurred. It takes them as reference for approximating their noise prominence of peaks. This step helps in determining the prominence threshold to later extract the spatial coordinates of local maxima in the rest of the images. The prominence 𝓅 approximation algorithm iterates over the noisy images, each iteration increments 𝓅_*n*_ value until the number of LM coordinates *l*_*n*_ = 0. Or when the incrementing iteration detects the same number of local maxima such as *l*_*n*_ = *l*_*n*-4_, then the program assumes that 𝓅 = 𝓅_*n*_ where “*n*” is the iteration number. These four images are also utilized to estimate the full width at half maximum (FWHM) *σ* over the noise LM, and to measure Δ*μ*, which is the average of the mean intensities Δ*μ*_*j*_ at LM *C*_*j*_ with radius *r* **Eq. (11)**

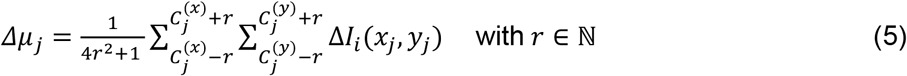

The region selection procedure is like that of parameter automation. We determine Δ*μ*_*j*_ and *σ*_*j*_ for each event *E*_*j*_ at *C*_*j*_. To designate *E*_*j*_ as a selected region we employ the following condition:

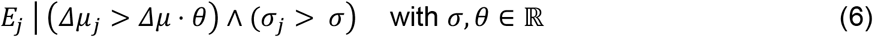

Here, θ denotes the sensitivity parameter, which can be adjusted by the user.

After detection, IVEA performs spatiotemporal tracking for recognition. This is applied for each selected region over a certain radius and period. Because the fixed FBE detection was used mainly to detect synaptic transmission in neurons, we implemented additional algorithm for detecting and tracking agonist/electric and 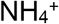 stimulations. This algorithm is important to recognize and sort events based on their occurrence period. Stimulus detection expressed as:

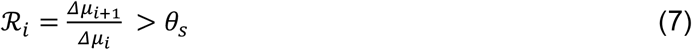

Where ℛ_*i*_ is the mean ratio, *θ*_s_ is the mean ratio threshold default is 1.1, Δ*μ*_*i*_ and Δ*μ*_*i*+l_ are the mean gray value of image Δ*I*_*i*_ and Δ*I*_*i*+l_ respectively.

To avoid an increase in the number of detected events due to high fluorescence intensity during the stimulus period, we adjust the detection sensitivity by increasing the detection sensitivity *θ*_*i*_, such as *θ*_*i*_ = *θ*⋅ ℛ_*i*_.

### Feature extraction

Feature extraction is performed by extracting sequence of image patches around *C*_*j*_ over a time interval. This patch is subdivided into smaller regions. Subsequently, we determine the mean intensity of each region. (**Fig. 2a**). For each selected region denoted as *E*_*j*_, we extract the spatial neighboring pixels around *C*_*j*_ as a 2D matrix ℳ^*j*^ ∈ ℝ^k×k^, where *k* is the window kernel defined by the user. The spatiotemporal data 𝒱_*j*_ ∈ ℝ^k×k×T^, which represents event *E*_*j*_ that occurred at time *t*_*j*_, is extracted over several frames T expressed as (3).

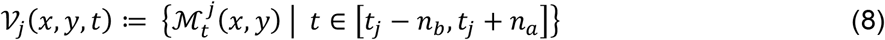

Whereas *n*_*b*_ is the number of frames before *t*_*j*_ and *n*_*a*_ is the number of frames after *t*_*j*_. Each matrix ℳ^*j*^ spatial coordinates are split into 13 small regions 𝔣 *ϵ* ℕ, which demonstrate the features matrix 𝒫_*j*_ *ϵ* ℝ^𝔣×T^ for event *E*_*j*_ such as (4):

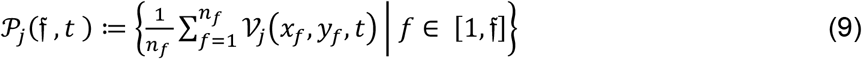

Whereas *n*_*f*_ is the number of pixels in each region *f*.

This approach, forms time series data 𝒫, represented as 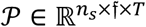, where *n*_s_ is the total number of nominated events. Each element 𝒫_*j*_ in 𝒫 represents 13 distinct signals that capture the fluorescence intensity profile of different regions plotted on a single graph (**Fig. 2b, Supplementary Fig. 5**). For fixed FBE we took *n*_*b*_ = 10 and *n*_*a*_ = 30 time samples, resulting in a total of *T* = 41 time samples. The LSTM model that was implemented for non-fixed FBE, the total number of time samples was *T* = 21, with *n*_*b*_ = 10 and *n*_*a*_ = 10. The time series data 𝒫 is first normalized within the range of 0 to 1 before being converted to tensors. For fixed FBE analysis, 𝒫 is converted into 3D tensors 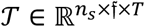 as follows:

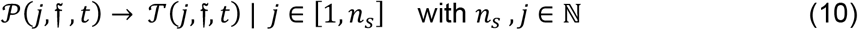

### Multivariate LSTM neural network architecture

Our LSTM network comprises four different layers. It serves as a robust framework for multivariate temporal data analysis. The first input layer, defined by the input shape specification, establishes the dimensions for the incoming multivariate time series data. This initial stage isn’t a distinct processing layer but rather a configuration step to align the network with the input data’s structure. The subsequent architecture unfolds with a convolutional 1D layer employing Rectified Linear Unit (ReLU). The subsequent layer incorporates a Long Short-Term Memory (LSTM) layer, designed to recognize sequential patterns. To promote stable training dynamics, batch-normalization layer is added. The last layer is the fully connected Dense layer employs the softmax activation function, rendering the architecture adept for multiclass classification.

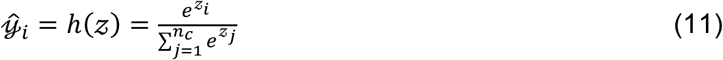

Whereas *h*(*z*) is the softmax function, *z*_*i*_ represents the raw score (logit) for a specific class *i* and *n*_*c*_ is the number of classes. This arrangement encapsulates both localized and temporal patterns inherent to multivariate sequential data, combining convolutional, recurrent, and normalization mechanisms. The network is structured to accommodate categorical cross-entropy as the loss function ℒ **Eq. (15)**, tailored for multiclass categorization, while optimization leverages the Adam optimizer with a learning rate 10^−3^.

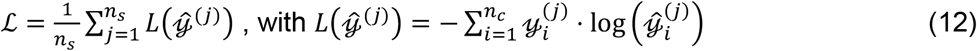

Where 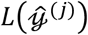 is single data point (sample), *n*_s_ is the number of data points, 𝓎_*i*_ ∈ {0, 1} is the true label for class *i* and 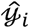 is the predicted probability.

### Encoder-ViT network architecture

Our eViT network consists of two components: a convolutional neural network (CNN) for feature extraction from image patches and a vision transformer (ViT) for classification. The encoder comprises seven layers, including a spatial convolution layer that is followed by a sequence of 3D convolutional layers and 3D max pooling operations (**Supplementary Fig. 11a,b**).

The encoder input layer accepts time series of 26 image patches with a width and height of 32×32, with the last dimension corresponding to the number of channels, such as 𝒳 ∈ ℝ^*t*×*w*×*h*×*c*^. If the dimension of the image patches changed, we use bilinear interpolation to resize the images. The initial stage of the encoder involves the application of a 3D convolutional layer with 8 filters and ReLU activation, which is followed by a 3D max pooling layer. Subsequently, another 3D convolutional layer with 16 filters with ReLU activation, followed by another 3D max pooling operation. Finally, a 3D convolutional layer with 32 filters and ReLU activation, followed by a concluding 3D max pooling layer. Each batch of the encoded image patches is passed through a Flatten layer and a Dense layer over time, then positional embedding is added to the data before passing them into a transformer block and subsequently into an MLP with a softmax activation for classification. The MLP is comprised of two Dense layers, the first layer has GeLU non-linearity.

### Neural network training

Both the LSTM and the eViT networks were trained using the Python programming language. The training data for the LSTM network was managed and prepared using the MATLAB platform for the visualization of image patch region patterns. In contrast, the eViT network training data was managed using ImageJ in the form of videos associated with their ROI files. These files contain the ROI’s center coordinates, frame position, and radius. Although our focus was on fusion events, we have selected other regions to train the neural network on various types of patterns, including motion patterns and certain types of noise that could potentially influence our neural network prediction. Using labeled data from HE and visualizing each event as a specific graph pattern, we were able to construct input training data samples for the LSTM and the eViT networks. For the LSTM the data was exported as CSV files the form of 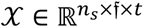 for fixed

FBEs, while the input training data category 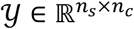, where, *n*_s_ is the number of samples and *n*_*c*_ is the number of classes. As for the eViT network, the data was saved as videos with their associated ROI files both zip and roi file container. The input data for the eViT network is 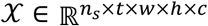 and 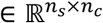. Our LSTM network for fixed FBEs was trained with 11.3k data samples, while for the non-fixed FBEs the LSTM network was trained on 12.6k data samples. The eViT network for non-fixed FBEs was trained with 548 videos approximately 7k data samples. These 548 videos were acquired at a rate of 10 Hz. For videos in which the eViT was tested and acquired at 50 Hz, the videos were reduced by a factor of five using the ImageJ “reduce” function, resulting in a rate of 10 Hz. We used Nvidia RTX 3070 for training our neural network, and Intel core i9-10920X CPU.

#### Training new data type

Training the neural network on new data involves two steps labeling the data and training the neural network. To label the data, the user can create ROI over the event using the ImageJ software and save them using the “ROI Manager” tool. Subsequently, the user should rename the ROI with a label and a number, then save the ROI/s as a roi/zip file under the same name as the video. The next step involves training the neural network using Python. Users should set up a Python development environment, such as the Anaconda platform. To initiate neural network training, users must first install the libraries associated with the Google TensorFlow platform. The script will prompt users to specify the directory containing their labeled data, including “videos and their ROIs”, as well as our prelabeled data files. Users can choose to combine their labeled data with our fusion events or only with the prelabeled noise and artifact data. The Python script generates a trained Keras model and saves it to the designated directory. IVEA enables users to import a custom model, which it will use for subsequent predictions. If no custom model is imported, IVEA will use the embedded neural network by default.

#### Google TensorFlow-Java implementation

IVEA LSTM and eViT network were developed using the Python v3.8.15 language and Google’s machine learning and artificial intelligence framework TensorFlow v2.9.1 or v2.10 with the Keras library. Using Python, we were able to train our neural network and export the trained model as a Protocol Buffer (pb) file format. To load and use our model with ImageJ Fiji, we used the Google TensorFlow Java v1.15.0 library and deeplearning4j core v1.0.0-M1.1 in our software. Integrating Google TensorFlow with Java is a complex task, particularly within the context of Fiji implementation. While Java offers versatility, it has limitations compared to Python, particularly in providing user-friendly and adaptable tools for machine learning applications. Notably, the Java support for Google TensorFlow is constrained, and as of the year 2024, faces issues with deprecated documentation. Additionally, the consolidation of all components into a single Java archive (jar) file poses challenges within the Fiji environment. In an effort to simplify the integration of the Google Deep Learning Framework with Java inside Fiji, we provided a concise explanation of the TensorFlow Java implementation on https://github.com/AbedChouaib/IVEA.

#### Video simulation and noise control

To mimic the CTL’s vesicles and simulate the fusion activity, we first create the vesicles as small round spheres with gaussian intensity spread using equation:

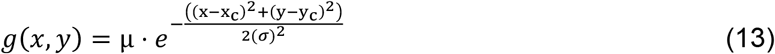

Whereas *μ* is the intensity of the spot and sigma the standard deviations controlling the spatial spread distribution.

We then added some random spatial movement for the vesicles to add motion variable. The vesicle’s fusion was more like fluorescence intensity cloud that spreads and disappears. To simulate these phenomena, we used Gaussian spread by time equation to control the temporal presence of the fusion by time, then we added one more variable for the radial spatial spreading dependent on the time variation. The overall equation expressed as:

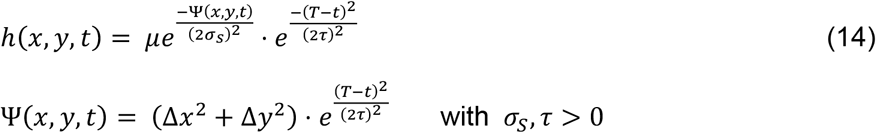

Whereas Ψ(*x, y, t*) simulate the radial dynamic dispersion of vesicle cargo over time, *t* is the current frame, *T* is the frame where the fusion occurs, *τ* is the fusion time interval, *σ*_s_ is the fusion radial spread and *μ* is the fluorescence intensity magnitude.

For the noise control analysis, we generated twenty distinct SNR levels between 10 and 0.1 SNR, simulating photon shot noise (Poisson noise) as commonly encountered in microscopy practices.

#### Hotspot area detection algorithm

The IVEA hotspot area extraction is based on DART algorithms, which employ unsupervised learning to segment the image into different layers. Following image segmentation, the MIC algorithm is performed to address the non-uniform regional fluctuations in fluorescence intensity, which is conducted prior to foreground subtraction. MIC is an enhanced version of the simple ratio and the previous method used with DART. MIC clusters the first image into a series of layers, wherein each layer comprises a group of labeled pixels that exhibit a close range of gray values, as determined by k-mean clustering. This process can be expressed as *I*(*x, y*) → *I*(*x, y, k*) | *k* = ℕ^5^ (**Supplementary Fig. 9**). Conventional approach (DART) involves the addition of the difference in gray values of clusters between two subsequent images. In contrast, with MIC we employed a simple ratio equation for each layer, assuming that the least cluster value represents the background. In the event of uniform regional fluctuations, the number of clusters of MIC could be reduced to k=1, this would yield to a result similar to that of the simple ratio. MIC is expressed as:

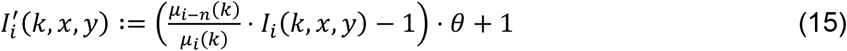

Where *i* is the *i*-th frame, *k* is the number of layers, *n* is the frame difference, and *θ* is a user input parameter added to control intensity adjustment, default *θ* = 1.

The iterative threshold consists of two distinct parts: initially, we capture two images where no events have occurred. Next, we compute Δ*I* and transform it into an 8-bit image to decrease computation time by reducing the iterations to under 255 steps. Finally, we attempt to clear Δ*I* repeatedly. The clearing processes consist of three sequential operations: threshold, erosion, and median filtering. In the iterative process, the threshold starts at half the mean intensity of Δ*I*, then we perform erosion with kernel *K*_e_ [*n, n*] to eliminate lone pixels Δ*I* = Δ*I* ⊖ *K*_*e*_ *as n* = 3. After erosion, a median filter with a user-defined radius or a preset default value is applied. The average mean gray value of the processed image is calculated and checked to see if it is equal to zero. If not, we iteratively increment the threshold by one gray step until we reach an average mean value of zero. The outcome of this process delivers the first iterative threshold decision *v*_l_. The second threshold decision *v*_2_ is performed for the remaining images, where this threshold is determined to correct the first threshold. The second threshold is like the previous process, except that a specific area of the segmented background is selected from each image (**Supplementary Fig. 8b**). The final threshold decision *v*_*i*_ is determined by *v*_*i*_ = *v*_2_ ⋅ *α* where *α* is the threshold sensitivity, if *α* was set as zero. The software takes two more frames to learn the sensitivity, it assumes no events had occurred and try to correct *v*_*i*_ by tuning *α* automatically. This step adjusts the difference between iterating over the entire image and iterating exclusively over the image’s background. Regions surpassing the global threshold *v*_*i*_ are considered as detected occurrences. Subsequently, each contiguous region is isolated and assigned a distinctive label designating it as an event. The fluorescence intensity of each event is spatially and temporally tracked immediately after detection. The mean intensity of each event *μ*_e_(*t*) is measured over time over a fixed area, then we calculate the mid intensity 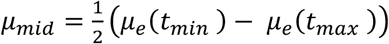 When the fluorescence intensity of the event falls below the mid intensity, the event signal is considered to have disappeared and the tracking stops (**Supplementary Fig. 8c,d**).

### Mice for T Cell, chromaffin cell and DRG neuron culture

WT mice with C57BL/6N background used in this study were purchased from Charles River. Synaptobrevin2-mRFP knock-in (KI) mice were generated as described in Matti *et al*.^13^. Granzyme B-mTFP KI mice with C57BL/6J background were generated as described previously^39^. Granzyme B-tdTomato KI mice^40^ were purchased from the Transgenesis and Archiving of Animal Models (TAAM) (National Centre of Scientific Research (CNRS), Orleans, France). Mice were housed in individually ventilated cages under specific pathogen-free conditions in a 12-h light-dark cycle with constant access to food and water. All experimental procedures were approved and performed according to the regulations by the state of Saarland (Landesamt für Verbraucherschutz, AZ.: 2.4.1.1).

### Murine CD8+ T cells

#### Culture

Splenocytes were isolated from 8–20-week-old mice of either sex as described before^4^. Briefly, naive CD8 T cells were positively isolated from splenocytes using Dynabeads FlowComp Mouse CD8+ kit (Invitrogen) as described by the manufacturer. The isolated naive CD8+ T cells were stimulated with anti-CD3ε /anti-CD28 activator beads (1:0.8 ratio) and cultured for 5 days at 37 °C with 5% CO_2_. Cells were cultured at a density of 1 × 10^6^ cells/ml in 12 well plates with AIMV medium (Invitrogen) containing 10% FCS, 50 U/ml penicillin, 50 *μ*g/ml streptomycin (Invitrogen), 30 U/ml recombinant IL-2 (Gibco) and 50 *μ*M 2-mercaptoethanol. Beads and IL-2 were removed from T cell culture 1 day before experiments.

#### Transfection and constructs

Day 4 effector T cells were transfected 12 h prior to the experiment through electroporation of the Plasmid DNA (Granzyme B-pHuji, Synaptobrevin2-pHuji) using Nucleofector™ 2b Device (Lonza) and the nucleofection kit for primary mouse T cells, according to the manufacturer’s protocol (Lonza). After nucleofection, cells were maintained in a recovery medium as described by Alawar *et al*. ^41^. 4 h prior to the experiment the cells were washed with AIMV medium (Invitrogen). The pMax_granzyme B-pHuji construct was generated by replacing the mTFP at the C-terminus of pMax-granzyme-mTFP^39^ with pHuji using a forward primer that included an AgeI restriction site 5′-ATG TAT ATC CAC CGG TCG CCA CCA TGG TGA GCA AGG GCG AGG AG-3′ and a reverse primer that included a NheI restriction site 5′-ATG TAT AGC TAG CTT ACT TGT ACA GCT C-3′. The size of this plasmid was 4.315 kb. Synaptobrevin2-pHuji plasmid was generated as described in ^42^.

#### Acquisition conditions

Measurement of exocytosis was performed via TIRFM as follows. We used day 5 bead activated CTLs isolated from GzmB-mTFP KI, GzmB-tdTomato KI, Synaptobrevin2-mRFP KI or WT mice. The latter were transfected with the above descripted constructs. 3 × 10^5^ cells were resuspended in 30 *μ*l of extracellular buffer (10 mM glucose, 5 mM HEPES, 155 mM NaCl, 4.5 mM KCl, and 2 mM MgCl_2_) and allowed to settle for 1–2 min on anti-CD3ε antibody (30 *μ*g/ml, BD Pharmingen, clone 145-2C11) coated coverslips. Cells were then perfused with extracellular buffer containing calcium (10 mM glucose, 5 mM HEPES, 140 mM NaCl, 4.5 mM KCl, 2 mM MgCl_2_ and 10 mM CaCl_2_) to stimulate CG secretion. Cells were recorded for 10 min at 20 ± 2 °C.

#### Imaging

Live cell imaging was done with two setups. The experiments performed with CTL (lytic granule staining with synaptobrevin-mRFP, granzyme B-mTFP or granzyme B-tdTomato) were performed with setup # 1 described previously^14, 32^. Briefly, an Olympus IX70 microscope (Olympus, Hamburg, Germany) was equipped with a 100x/1.45 NA Plan Apochromat Olympus objective, a TILL-total internal reflection fluorescence (TILL-TIRF) condenser (TILL Photonics, Kaufbeuren, Germany), and a QuantEM 512SC camera (Photometrics) or Prime 95 B scientific CMOS camera (Teledyne Photometrics, Tucson, AZ, USA). The final pixel size was 160 nm and 110 nm, respectively. A multi-band argon laser (Spectra-Physics, Stahnsdorf, Germany) emitting at 488 nm was used to excite SypHy and mTFP fluorescence, and a solid-state laser 85 YCA emitting at 561 nm (Melles Griot Laser Group, Carlsbad, CA, USA) was used to excite mRFP and tdTomato. The setup was controlled by Visiview software (Version:4.0.0.11, Visitron GmbH). The acquisition frequency was 10 Hz for all experiments.

The setup # 2 used to acquire CTL secretion, in which the lytic granules were labeled by Synaptobrevin-pHuji and granzyme B-pHuji, was previously described^39, 42^. Briefly the setup from Visitron Systems GmbH (Puchheim, Germany) was based on an IX83 (Olympus) equipped with the Olympus autofocus module, a UAPON100XOTIRF NA 1.49 objective (Olympus), a 445 nm laser (100 mW), a 488 nm laser (100 mW) and a solid-state 561 nm laser (100 mW). The TIRFM angle was controlled by the iLAS2 illumination control system (Roper Scientific SAS, France). Images were acquired with a QuantEM 512SC camera (Photometrics) or Prime 95 B scientific CMOS camera (Teledyne). The final pixel size was 160 nm and 110 nm, respectively. The setup was controlled by Visiview software (Version:4.0.0.11, Visitron GmbH). The acquisition frequency was 5 or 10 Hz, and the acquisition time was 10 to 15 min.

### Murine DRG neurons

#### Culture and transfection

The training of the Fixed FBE neural network and the automatic detection of neuronal exocytosis at synapse was performed on data sets that were previously published^32, 33^. Shortly DRG neuron cultures from young adult (1–4 weeks old) WT of either sex was made as previously described^14^. Lentivirus infection to transfect with SypHy was performed on DIV1. The following day, the lentivirus was removed by washing before adding the second order spinal cord (SC) interneurons (SC neurons) to the culture to allow DRG neurons to form synapses. SC neurons were prepared from WT P0-P2 pups of either sex using as previously described^32^. DRG/SC co-culture was maintained in Neurobasal A (NBA) medium (Thermo Fisher Scientific, Waltham, MA, USA) supplemented with fetal calf serum (5% v/v), penicillin and streptomycin (0.2% each), B27 supplement (2%), GlutaMAX (1%, all Thermo Fisher Scientific, Waltham, MA, USA), and human beta-nerve growth factor (0.2 µg/mL, Alomone Labs, Jerusalem, Israel) at 37 °C and 5% CO2.

#### Acquisition conditions

Secretion was evoked by electrical stimulation via a bipolar platinum-iridium field electrode (#PI2ST30.5B10, MicroProbes, Gaithersburg, MD, USA) and a pulse stimulator (Isolated Pulse Stimulator Model 2100, A-M Systems, Sequim, WA, USA). The measurement protocol was 30 seconds without stimulus followed by a biphasic 1 ms long 4 V stimulus train at 10 Hz for 30 seconds to elicit exocytosis of SVs. At the end of the measurement, NH_4_Cl was applied to visualize the entire SV pool. During the measurement, the temperature was maintained at 32°C by a perfusion system with an inline solution heater (Warner Instruments, Holliston, MA, USA). The extracellular solution contained 147 mM NaCl, 2.4 mM KCl, 2.5 mM CaCl_2_, 1.2 mM MgCl_2_, 10 mM HEPES, and 10 mM glucose (pH 7.4; 300 mOsm). The NH_4_Cl solution had the same composition as the extracellular solution, but the NaCl was replaced with 40 mM NH4Cl.

#### Imaging

All experiments were performed on Setup # 1 described above for the CTLs.

### Chromaffin cells

Data showing bovine chromaffin cells exocytosis was from Becherer *et al*.^43^ and Hugo *et al*.^13^. Culture condition was described by Ashery *et al*.^44^. Chromaffin cells were electroporated with NPY-mRFP to label the large dense core vesicles. Secretion was induced through either depolarization trains^43^ or perfusion of the cells with 5 *μ*M Ca^2+^ containing solution via the patch-clamp pipette^13^. The acquisition rate was 10 Hz and the exposure time was 100 ms. The camera was either a Micromax 512BFT camera (Princeton Instruments Inc., Trenton, NJ, USA) with 100×/1.45 NA Plan Apochromat Olympus objective^43^, or a QuantEM 512SC camera (photometrics) with an 100×/1.45 NA Fluar (Zeiss) objective ^13^, giving a final pixel size of 130 or 116 nm^2^ respectively.

### INS-1 cells

#### Culture and transfection

Rat insulinoma cells^45^ (INS-1 cells, clone 832/13 provided by Hendrik Mulder, Lund University) were maintained in RPMI 1640 (Invitrogen) containing 10mM glucose and supplemented with 10% fetal bovine serum, streptomycin (100 µg ml^-1^), penicillin (100 µg ml-1), Na-pyruvate (1 mM) and 2-mercaptoethanol (50 µM). The cells were plated on polylysine-coated coverslips, transfected using lipofectamine 2000 (Invitrogen), and imaged 24-42 h later.

#### Acquisition conditions

The bath solution contained (in mM) 138 NaCl, 5.6 KCl, 1.2 MgCl_2_, 2.6 CaCl_2_, 10 D-glucose, 0.2 Diazoxide, 0.2 forskolin, and 10 HEPES, pH 7.4 adjusted with NaOH. Individual cells were stimulated by computer-controlled air pressure ejection of a solution containing elevated K^+^ (75 mM replacing Na^+^) through a pulled glass pipette (similar to patch clamp electrode) that was placed near the recorded cell. The bath solution temperature was kept at 35°C.

#### Imaging

INS1 cells (clone 832/13) that transiently expressed NPY-mGFP, NPY-mNeonGreen or NPY-mCherry were imaged using a custom-built lens-type total internal reflection (TIRF) microscopes based on AxioObserver D1 microscope with an x100/1.46 objective (Carl Zeiss). Excitation was from a diode laser module at 473 nm, or a diode pumped laser at 561 nm, respectively (Cobolt, Göteborg, Sweden), controlled by an acoustic-optical tunable filter (AOTF, AA-Opto, France). Light passed through a dichroic Di01-R488/561 (Semrock), and emission light was separated onto the two halves of a sCMOS camera (Prime 95B, Teledyne photometrics) using an image splitter (Dual view, Photometrics) with a cutoff at 565 nm (565dcxr, Chroma) and emission filters (FF01-523/610, Semrock; and ET525/50m and 600EFLP, both from Chroma). Scaling was 110 nm per pixel (sCMOS camera). The acquisition rate for NPY-mNeonGreen was 50 Hz and NPY-mCherry 10Hz. NPY-eGFP expressing INS1 cells were imaged using a TIRF microscope that was based on an AxioObserver Z1 (Zeiss) with a diode pumped laser at 491 nm (Cobolt, Stockholm, Sweden) that passed through a cleanup filter and dicroic filter set (zet405/488/561/640x, Chroma). Imaging was done with a 16-bit EMCCD camera (QuantEM 512SC, Roper) with a final scale of 160 nm per pixel. The acquisition rate was 10 Hz. Image acquisition was conducted with MetaMorph (Molecuar Devices).

### Human CD8+ T lymphocytes

*Cells*. Human CD8+ T cell clones were used as cellular model. Human T cell clones were isolated and maintained as previously described (Filali *et al*. 2022). Briefly, cells were cultured in RPMI

1640 medium GlutaMAX (Invitrogen) supplemented with 5% human AB serum (Institut de Biotechnologies Jacques Boy), 50 *μ*M 2-mercaptoethanol, 10 mM Hepes, 1× MEM-Non-Essential Amino Acids (MEM-NEAA) Solution (Gibco), 1× sodium pyruvate (Sigma-Aldrich), ciprofloxacin (10 *μ*g/ml) (AppliChem), human recombinant interleukin-2 (rIL-2; 100 IU/ml), and human rIL-15 (50 ng/ml) (Miltenyi Biotec). Blood samples were collected and processed following standard ethical procedures after obtaining written informed consent from each donor and approval by the French Ministry of the Research as described (Cortacero *et al*. 2023, authorization no. DC-2021-4673).

#### Acquisition conditions

Human CTLs were stained for 30 min with Lysotracker red DND-99 (2µM) (Invitrogen) at 37°C/5% CO_2_. The cells were washed 3 times with RPMI 1640 medium (1X) w/o pH Red supplemented with 10 mM L-Glutamine, 10mM of Hepes. µ-Slide 15 Well 3D glass bottom slides (Ibidi) were coated with poly-D-lysine (Sigma), human monoclonal anti-CD3 antibody (5µg/mL or 10µg/mL) (TR66) (Enzo Life Sciences) and recombinant human ICAM-1/CD54 Fc Chimera Protein (5µg/mL or 10µg/mL) (R&D Systems) at 4°C overnight. The chambers slides were washed 3 times with PBS and mounted on a heated stage within a temperature-controlled chamber maintained at 37°C and constant 5% CO_2_. For each recording, 3 × 10^4^ to 5 × 10^4^ cells were seeded on the chambered slides. During acquisition, the cells were in RPMI 1640 medium (1X) w/o pH Red supplemented with 10 mM L-Glutamine, 10mM of Hepes and 5% fetal calf serum (FCS).

#### Imaging

The TIRFM set up acquisition was based on an Eclipse Ti2-E inverted microscope (Nikon) equipped with a 100×/1.45 NA Plan Apochromat LBDA objective (Nikon Instruments) and an iLAS 2 illumination control system (Roper Scientific SAS). A diode laser at 561nm (150 mW) (Coherent) band-passed using a ZET405/488/561/647x filter (Chroma Technology) was used for excitation. The emissions were separated using a ZT405/488/561/647rpc-UF1 dichroic (Chroma Technology) and optically filtered using ZET405/488/561/647m filter (Chroma Technology). Images were recorded on a Prime 95B Scientific CMOS Camera (Teledyne Photometrics). The final pixel size was 110 nm. Image acquisition was controlled using MetaMorph Software (Version 7.10.5.476, Molecular Devices) and Modular V2.0 GATACA software. The acquisition frequency was 9 Hz for a duration of 20 to 30 minutes.

### Dopaminergic neurons

Data showing dopaminergic neuron exocytosis monitored by AndromeDA nanosensor paint technology were from Elizarova *et al*.^24^. The setup included a 100× oil-immersion objective (UPLSAPO100XS, Olympus) and a Xenics Cheetah-640-TE1 InGaAs camera (Xenics) giving a final pixel size of 150 nm. The imaging was performed at 15 Hz.

### Statistical analysis

All statistical analyses were performed with SigmaPlot (V14.5.0.101, Systat Software, Inc.). P-values were calculated with two-tailed statistical tests and 95% confidence intervals. ANOVA, ANOVA on ranks and Student’s t-test were used as required (see **Supplementary Fig. 1**).

## Ethics statement

Mice were treated according to the regulations of the local authorities, the state of Saarland (Landesamt für Verbraucherschutz) under the license AZ.: 2.4.1.1 or the Niedersächsisches Landesamt für Verbraucherschutz und Lebensmittelsicherheit (LAVES, permit numbers 33.19-42502-04-19/3254, 33.19.42502-04-15/1817 and 33.19-42502-04-18/2756). Animals were housed according to European Union Directive 63/2010/EU and ETS 123 at 21 +/-1°C, 55% relative humidity, under a 12 h/12 h light/dark cycle, and received food and tap water ad libitum. Human blood samples were collected and processed following standard ethical procedures after obtaining written informed consent from each donor and approval by the French Ministry of the Research (authorization no. DC-2021-4673).

## Data availability

All original data sets used for this study and presented in the Source Data file will be deposited in ZENODO repository server (https://zenodo.org/) upon acceptance. The accession code will be provided by the corresponding author upon reasonable request. The program and the source code are available on GitHub (https://github.com/AbedChouaib/IVEA).

## Acknowledgments

This work was supported by grants from the Deutsche Forschungsgemeinschaft (SFB 894 to U.B.), the European Commission (ERC-2021-SyG_951329 to the Department of Cellular Neurophysiology, Saarland University and to S.V., Cancer Research Center of Toulouse, INSERM UMR1037) and grants from University of Saarland (HOMFORexzellent2020 and NanoBioMed Young Investigator grant 2020 to H.-F.C.). A.H.S. was funded by the European Research Council (ERC) under the European Union’s Horizon 2020 research and innovation program (grant agreement No 951275). C.P. was supported by the Deutsche Forschungsgemeinschaft (DFG, German Research Foundation) under Germany’s Excellence Strategy - EXC 2067/1-390729940. We thank Hendrik Mulder, Lund University for providing the INS-1 cells. We thank Marcel Lauterbach, Eugenio F. Fornasierio and Silvio O. Rizzoli for their constructive comments on the initial version of the manuscript. We thank Margarete Klose, Anja Bergsträßer, Nicole Rothgerber for excellent technical assistance.

## Author Contributions

A.H.S. and U.B. conceived and supervised the project. A.A.C. developed the methodology, investigated and analyzed the data. C.P. provided critical input on the developed artificial intelligence strategies. A.A.C., U.B, H.-F.C., O.M.K, L.D., S.Ec. were the human experts. H.-F.C., O.M.K., S.Ec., L.D., S.El., J.A.D., A.H.S. and U.B. acquired the original biological data. U.B. acquired and managed the project funding. Additional funding was acquired by S.V., S.B. and C.P. A.A.C, A.H.S and U.B. wrote the manuscript which was revised by all authors.

## Competing interests

The authors declare no competing interests.

## Notes

https://github.com/AbedChouaib/IVEA

### Competing Interest Statement

The authors have declared no competing interest.

