## Supplementary information for "Highly adaptable deep-learning platform for automated detection and analysis of vesicle exocytosis"

### Supplementary Figures

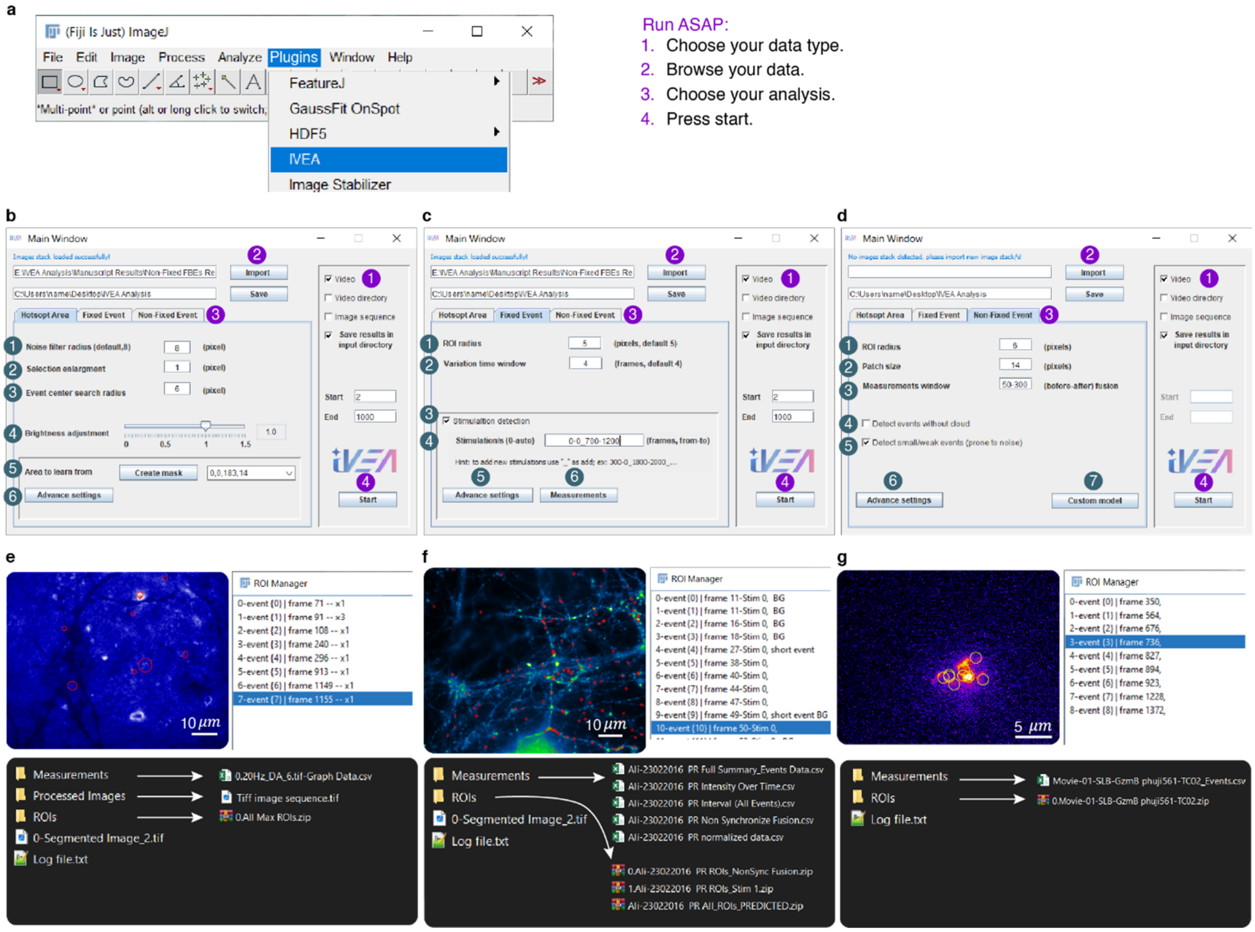

**Supplementary Figure 1. IVEA GUI. a**, Plugin location in ImageJ Fiji. **b**, Hotspot Area Events Analysis: Noise filter used for extracting events from the image, the higher it goes the less sensitive detection will be (This filter doesn't affect the measurements). 2- Enlarge the mask by n pixels. 3- Search radius to identifying the events identity. 4- Brightness adjustment sensitivity (used in case of image intensity fluctuation correction). 5- Select area for the program to learn from. 6- Advance settings containing control parameters that are mostly determined automatically by the software. **c**, Fixed FBEs Analysis: 1- Size of the ROI radius around an event. 2- Adjusts the time frame for foreground detection variation. 3- Enables stimulus detection if checked, otherwise simulation will be excluded. 4- Simulation timing from a to b (a-b). a or b are determined automatically if set to 0. 5- Advanced settings for fixed FBE default parameters. 6- Measurement settings for output data. **d**, Non-Fixed FBEs Analysis: 1- Size of the ROI radius around an event. 2- Sequence image patch size. 3- Measurement time interval from a to b (a-b). 4- Include pH-insensitive events. 5- Include small/weak events. 6- Advanced settings for non-fixed FBE default parameters. 7- Controls the export of training data and the use of custom neural networks. **e**, Hotspot Area Events Analysis output. **f**, Fixed FBEs Analysis output. **g**, Non-Fixed FBEs Analysis output.

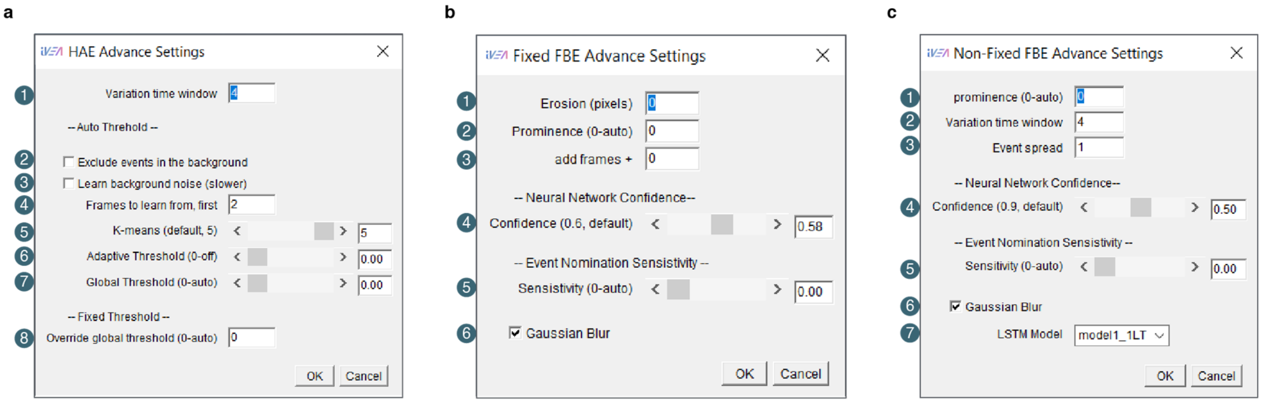

**Supplementary Figure 2. IVEA advance settings GUI. a**, Hotspot Area Events advance settings: time frame variation interval. 2- Exclude events that occur in the background. 3- Learn noise from the background. 4- Specify the frame used for IVEA to learn the threshold from. 5- Number of layers for image segmentation. 6- Adaptive threshold used to locally threshold hotspot areas and extract brightest spots. 7- Threshold sensitivity is automatically detected, 8- Override global threshold. **b**, Fixed FBEs advance settings: **1-** Erosion used only if user want to eliminate small events. 2- **"**Find Maxima" prominence. 3- Add frames to analyze by the neural network. 4- Neural network confidence level. 5- Detection sensitivity. 6- Gaussian blur filter. **c**, Non-Fixed FBEs advance settings: 1- "Find Maxima" prominence. 2- time frame variation interval. 3- Increasing the sigma of the event's spread in the medium. 4- Neural network confidence level. 5- Detection sensitivity. 6- Gaussian blur filter.

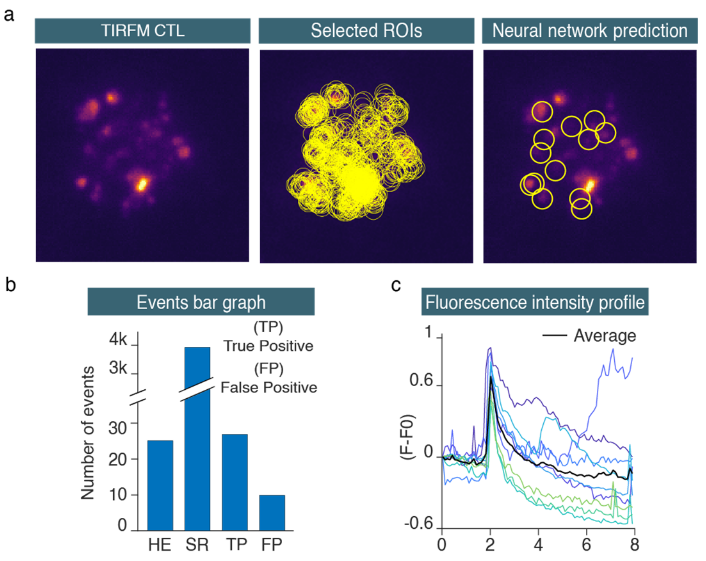

**Supplementary Figure 3. CTL analysis – Exocytosis of Lysotracker red labeled lytic granules. a**, The first image shows the raw TIRFM image of a CTL cell in which the lytic granules were labeled with Lysotracker red. The second image is the overlay of the selected regions of interest prior to importation into the neural network. The final image displays the overlay of the neural network prediction, indicating the events that were predicted to be true. **b**, The bar graph illustrates the comparative outcomes between the human expert (HE) and the IVEA software. The selected ROIs are indicated by "SR." **c**, The graph shows the intensity profile of the fusion events detected as true positive. Images acquired at the Cancer Research Center of Toulouse, France.

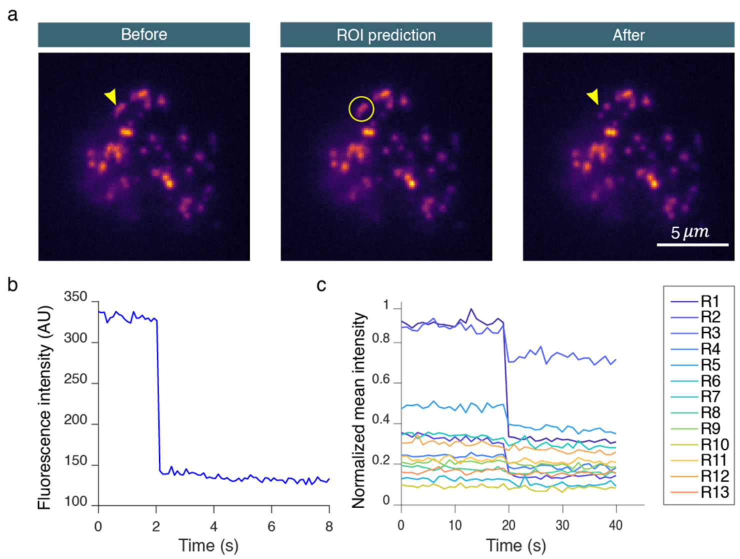

**Supplementary Figure 4. Chromaffin cell analysis. a**, Images of a chromaffin cell showing an exocytotic event detected by IVEA. Granules are stained through over-expression of NPY-mCherry (pH-insensitive, data from[1]). **b**, Fluorescence intensity profile of the event over 80 frames. **c**, Exocytosis pattern is different from another pattern the LSTM network was trained on. This data was better analyzed with the encoder-ViT network.

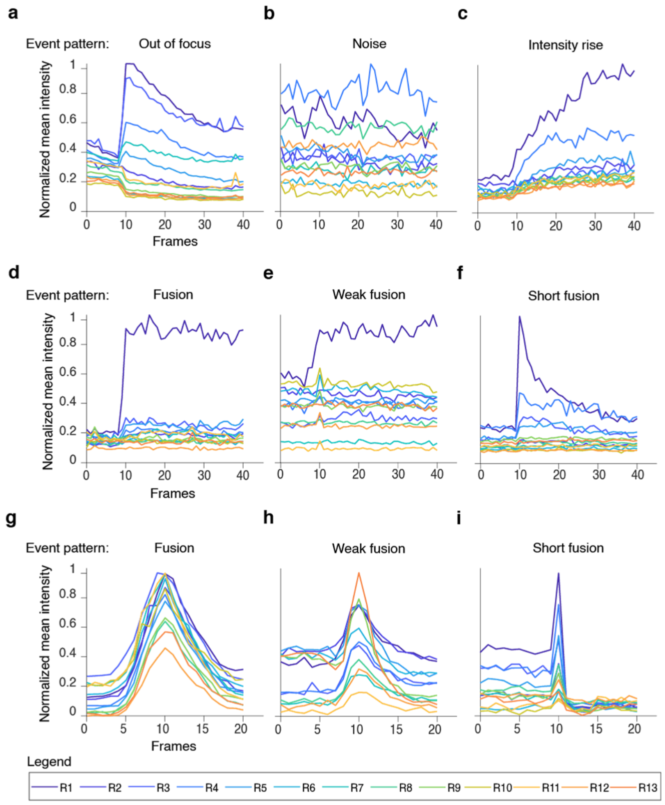

**Supplementary Figure 5. Events' patterns for LSTM network. a**, Out of focus patterns for DRG neurons. **b**, Random noise. **c**, intensity rise. **d**, DRG neurons fusion event. **e**, DRG neurons weak fusion event. **f**, DRG neurons short fusion event. **g**, T cell fusion event. **h**, T cell weak fusion event pattern. **i**, T cell short fusion event pattern.

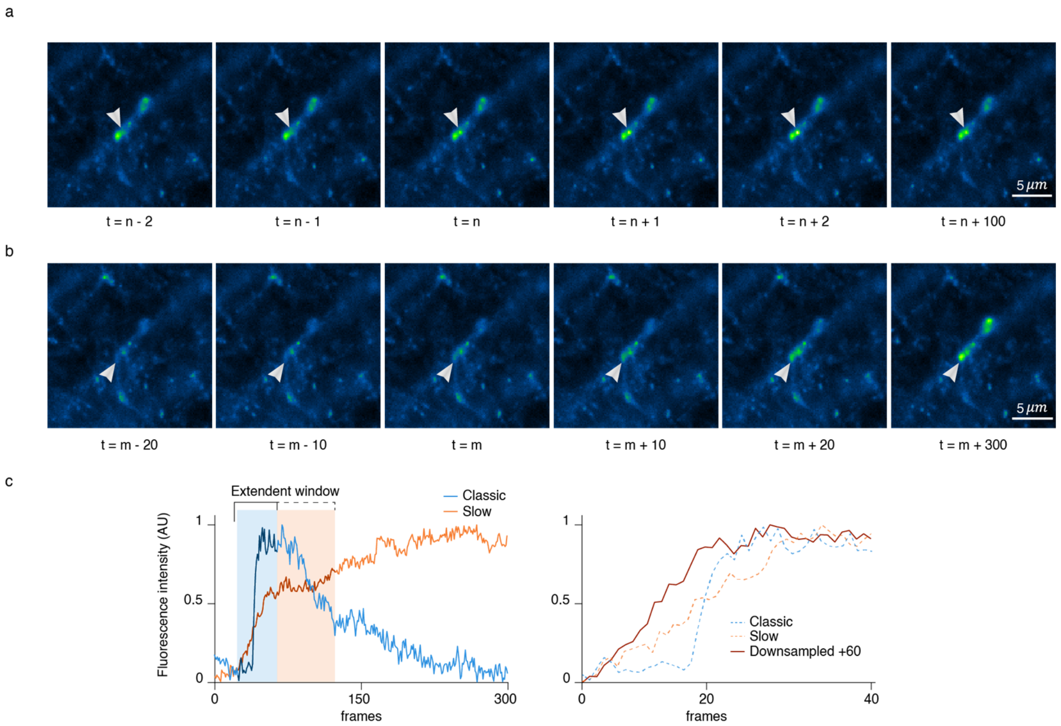

**Supplementary Figure 6. Slow vs classic fusion events. a**, Represents the timeline of classic fusion event. **b**, represent the timeline of slow fusion event. **c**, The first graph represents the event intensity profiles over 300 frames. The blue shade area is the default measurement window, while the orange shade area is the extended window adjusted by the user. The second graph compares the down sampled signal with the classic fusion event and the rise intensity event.

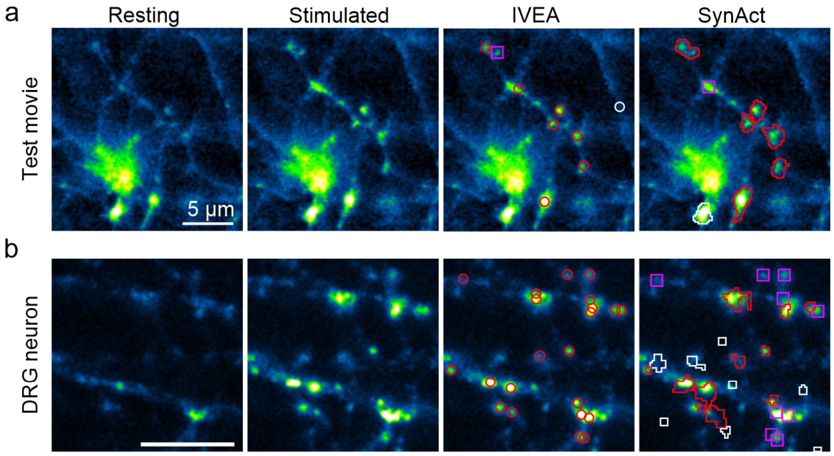

c

| SynActJ Settings | | Test movie | DRG neuron |
|---|---|---|---|
| Background | Gauss sigma | 4 | 4 |
| | Selected threshold | MinError | MinError |
| Boutons : Detect | LoG sigma | 0.5 | 0.5 |
| | Prominence | 0.005 | 0.005 |
| Segment | Gauss sigma | 1.0 | 0.5 |
| | RollingBall radius | 30 | 20 |
| | Selected threshold | Triangle | IJ -IsoData |
| | Erode Mask | OFF | ON |

**Supplementary Figure 7. IVEA is superior to SynActJ in detecting active synapses. a-b**, comparison of IVEA and SynActJ for the detection of active synapses. From left right are shown an image of the neurons at rest, during stimulation, with the labelled synapses detected by IVEA and the detected synapses by SynActJ. Shown in red are the overlayed ROIs of the true positive events, in white are the false positive events and the magenta squares are the missed events. **a**, Test movie provided with SynActJ. SynActJ settings were default values as recommended. For IVEA analysis, due to the small sequence size, the first and the last frames of the movie had to be duplicated so that the sequence reached 60 frames in length. **b**, Movie acquired with DRG-neurons over-expressing SypHy. For this analysis the movie was restricted to 300 frames encompassing the stimulation at frame 60. The raw data was analyzed by IVEA whereas frame reduction and a gaussian blur of 0.5 had to be applied to enable SynActJ to detect any event. SynActJ settings where adjusted iteratively to improve detection. **c**, SynActJ settings used to analyze the movies shown in ***a*** and ***b***. Default settings were used for IVEA's analysis for the movie shown in (**b**), while for the movie shown in (**a**), the automatic parameters were set off, the prominence was set to 10 and the sensitivity to 1.

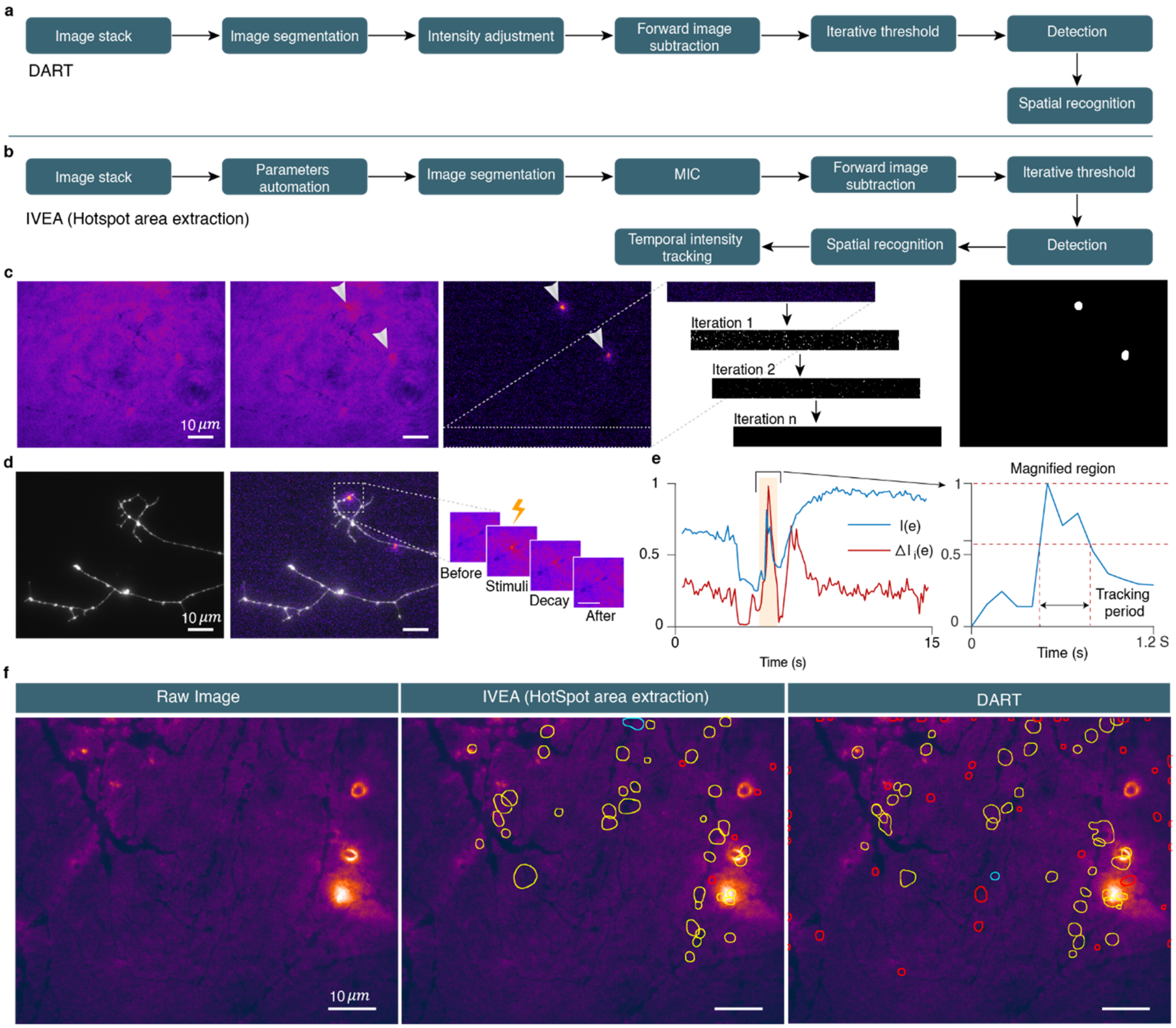

**Supplementary Figure 8. Overview sensor-based exocytosis detection algorithm. a**, Algorithm flowchart of DART. **b**, Algorithm flowchart of IVEA hotspots area extraction. IVEA employs three new methods in comparison to DART. These include parameter automation, multilayer intensity correction (MIC) (see **Supplementary Fig. 9**), and temporal tracking. **c**, Detection process from left to right: first is the raw image; second is the hotspot image with two hotspots in this frame; third is the intensity variation image where hotspots are clearly visible. The iteration threshold in this image was taking cropped area from which it learned the noise level by iterating and eliminating the noise; last is the segmented image that is used to determine the event's ROIs. **d**, Intensity tracking process is shown from left to right in three images. The first image displays the raw image of a neuron, the second image shows the corresponding intensity variation with the raw image overlaid, and the third image is a sequence of snapshots that display the hotspot over time delineated on the second image. **e**, Graph representing the intensity variation overtime for ROIs mean intensity of the original image sequence (I(e)) and for the intensity variation processed image sequence ($\Delta I_i(e)$). The graph on the right magnifies I(e) for the time window of the hotspot occurrence. It displays the temporal intensity tracking period. **f**, Images representing the comparative results between IVEA hotspot area extraction and DART. The yellow ROIs are the hotspot areas, the red ROIs are most probably false hotspots, and the cyan ROIs are true hotspots detected by one algorithm but not the other.

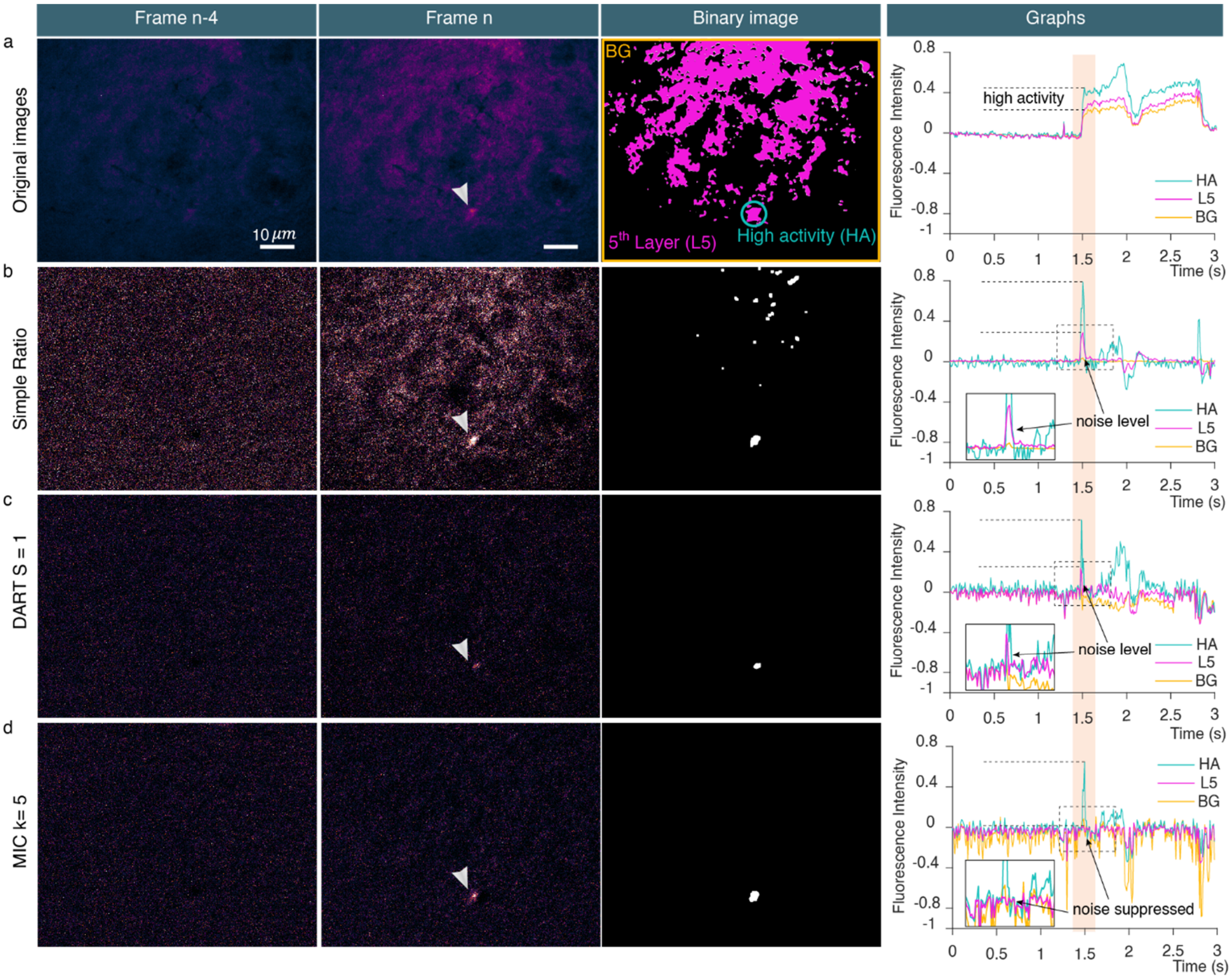

**Supplementary Figure 9. Multi-layer intensity correction (MIC).** The figure displays the results of three different algorithms, which have been employed to address non-uniform regional fluctuations in fluorescence intensity in videos. The first column represents an image without fluctuations. The middle column corresponds to an image containing a region of high activity and a background with fluorescence intensity fluctuation. The three last images in the third column show the binary images of the second column. They were generated by applying a triangular threshold, followed by morphological reconstruction comprising erosion, a median filter, and then dilation. The last column represents the fluorescence intensity profiles across three distinct regions that are displayed in the first binary image. These include the fifth segmented layer highest k-means cluster value (L5), represented in pink; the high activity region (HA), shown in blue; and the background (BG), represented in orange. The inset is a magnification of the curves delineated by a stippled line. **a**, Shows the raw images, while the binary image represents the L5 of the raw clustered image using k-means (k=5). The graph shows the high activity of the HA region compared to the L5 and BG regions over time. **b**, Impact of using simple ratio on non-uniform fluctuations in the image. The binary image shows how simple ratio failed to eliminate the false regions. The graph illustrates the presence of remaining noise in the measurement of the HA region. **c**, This panel shows the results of the previous method DART. Note that it partially addresses the non-uniform fluctuations. **d**, MIC (k=5) was able to suppress the noise entirely while preserving the HA.

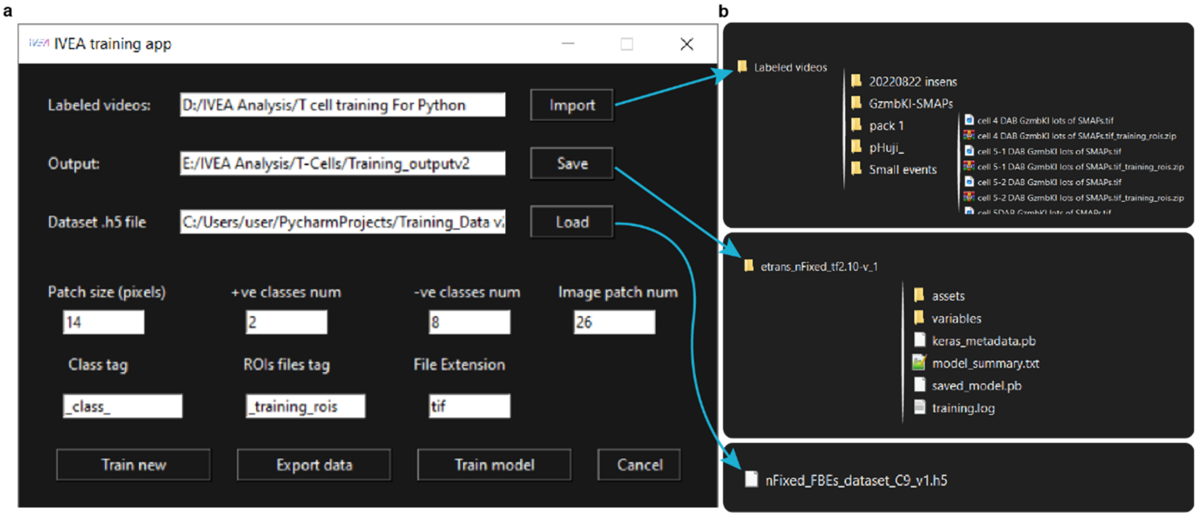

**Supplementary Figure 10. IVEA training GUI. a,** This panel represents the main interface for IVEA's training application (app). The "Import" button is utilized for the loading of the user's labeled videos directory. The "Save" button is used to store the trained model in the specified directory. The "Load" button is used to load a prelabeled dataset ".5h" file. The field "patch size" refers to the radius of the neural network image patch. The value entered in the field "+ve classes num" represents the number of classes that the user wishes to include. The number of classes to be excluded (i.e., not representing an exocytosis event) is indicated by "-ve classes num." The field "image patch num" refers to the number of time series images that are imported into the neural network. The field "class tag" is used to point the app at the class label. The "ROIs file tag" is a designation for a region of interest (ROI) file, which the app will search for in the suffix of the video name. The "File extension" field indicates the file type of the imported videos. The "Train new" button is employed when the user wishes to commence training from scratch. The "Export data" button is used to combine a new training set with the existing training dataset and export the new dataset as a ".h5" file. **b,** Exemplary representation of the input and output directories and files. The IVEA training app is capable of automatically searching subfolders and retrieving the associated videos and their respective ROI files.

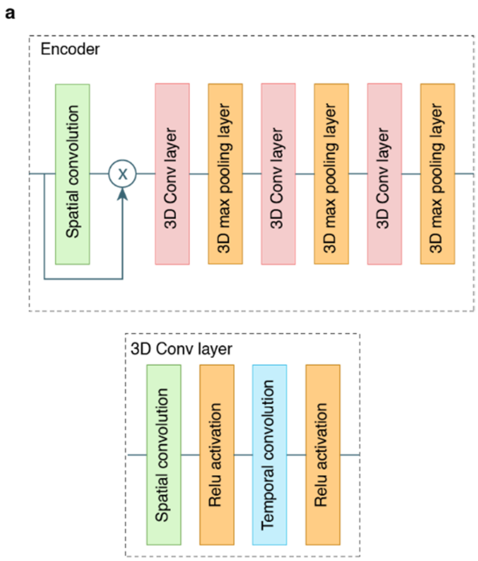

**Supplementary Figure 11. Encoder architecture. a**, Encoder layers, spatial convolution followed by 3D conv layer 3D max pooling. **b**, 3D conv layer consisting of separate spatial convolution followed by temporal convolution along the time axis followed by ReLU activation.

1. Becherer, U. et al. Quantifying exocytosis by combination of membrane capacitance measurements and total internal reflection fluorescence microscopy in chromaffin cells. *PLoS One* **2**, e505 (2007).

### Supplementary Tables

**Supplementary Table 1 (related to Fig. 4): statistical comparison of our eViT with the LSM and ExoJ.** Given are the p value of the performed tests. ANOVA on rank with Dunn's post-test is marked with an *, ANOVA with Holm-Sidak post-test is marked with # and Student two tailed paired t-test is marked with §. Non-significant differences are shown as ns. The p value is indicated in brackets when it is below 0.2. Pairs that could not be tested due to many missing values, are labeled with //.

| | eViT to LSTM | eViT to ExoJ | LSTM to ExoJ | N |
|---|---|---|---|---|
| Recall * | ns | <0.001 | 0.011 | 10 |
| Precision * | ns | ns | ns (0.134) | 10 |
| F1 score * | ns | <0.001 | 0.006 | 10 |
| Recall * | ns | <0.001 | 0.020 | 7 |
| Precision # | ns (0.155) | <0.001 | <0.001 | 7 |
| F1 score * | ns | <0.001 | <0.001 | 7 |
| Recall * | 0.033 | <0.001 | 0.033 | 10 |
| Precision * | ns (0.126) | 0.016 | <0.001 | 10 |
| F1 score § | 0.002 | // | // | 10 |
| Recall * | 0.030 | <0.001 | ns | 8 |
| Precision # | ns | <0.001 | 0.002 | 8 |
| F1 score § | 0.00000793 | // | // | 8 |

**Supplementary Table 2 (related to Fig. 4): Iterative adjustment of ExoJ analysis parameter.** Ten different parameters can be adjusted in ExoJ to improve event detection. We found that the first 8 parameters (Min points for fitting procedure down to Max. displacement) did not alter event detection. In contrast the last two parameters (Min. $R^2$ (Decay fitting procedure, DFP) and Min. $R^2$ (Est. radius fitting procedure, RFP) greatly affected event detection. We started the iterative adjustment with the default value then we first adjusted RFP and then DFP until the best possible detection was achieved. Metrics are: true positive (TP), false positive (FP) and false negative (FN) events. Test was performed on one movie from **Fig 4a**.

| Metric | ExoJ | | | | | | | eViT |
|---|---|---|---|---|---|---|---|---|
| | Default<br>0.9 – 0.75 | DFP – RFP<br>0.9 – **0.4** | DFP – RFP<br>0.9 – **0.5** | DFP – RFP<br>0.9 – **0.6** | DFP – RFP<br>**0.8** – 0.5 | DFP – RFP<br>**0.9** – 0.5 | DFP – RFP<br>**0.95** – 0.5 | Default |
| TP | 3 | 5 | 5 | 3 | 5 | 5 | 2 | 14 |
| FP | 1 | 11 | 5 | 1 | 9 | 5 | 1 | 2 |
| FN | 11 | 9 | 9 | 11 | 9 | 9 | 12 | 0 |

**Supplementary Table 3:** Comparison of our eViT with pHusion. The results of both programs are shown for the movies of each CTL labeled with the pH sensitive GzmB-pHuji (**Fig. 4a**). Movies of cells that did not yield any result using pHusion are labeled with //. We adjusted the Grid spacing to 0.11 µm (pixel size) and in the "good trace" selection we used an R2 function value of 0.9. All other pHusion parameters were default parameters. Displayed are the result from "traces collapsed to single event". Shown are the true positive (TP), false positive (FP) and false negative (FN) events.

| | pHusion software | | | IVEA (eViT) | | | HE |
|---|---|---|---|---|---|---|---|
| **Cells** | TP | FP | FN | TP | FP | FN | TP |
| **1** | // | // | // | 4 | 0 | 0 | 4 |
| **2** | // | // | // | 4 | 0 | 0 | 4 |
| **3** | // | // | // | 4 | 0 | 0 | 3 |
| **4** | // | // | // | 2 | 0 | 0 | 2 |
| **5** | 0 | 0 | 16 | 16 | 0 | 0 | 14 |
| **6** | // | // | // | 10 | 1 | 0 | 7 |
| **7** | 1 | 3 | 3 | 4 | 0 | 0 | 2 |
| **8** | 14 | 2 | 0 | 14 | 2 | 0 | 14 |
| **9** | 6 | 3 | 2 | 8 | 1 | 0 | 8 |
| **10** | 4 | 3 | 3 | 7 | 0 | 0 | 7 |
| **Total** | 25 | 11 | 24 | 73 | 4 | 0 | 65 |

### Supplementary Videos

**Supplementary Video 1. Exemplary video of an exocytosis event of a lytic granule visualized through pH-sensitive fluorescent cargo protein.** Cytotoxic T cell secretion was stimulated by seeding the cells on anti-CD3ε antibody. Granules are stained through over expression of pH-sensitive granzyme B-pHuji. Movies were acquired at 10 Hz; video display speed is ×2. The exocytotic events is displayed on the left. On the right is the fluorescence intensity graph for each event aligned at the detected frame.

**Supplementary Video 2. Exemplary video of an exocytosis event of a lytic granule visualized with pH-insensitive fluorescent cargo protein.** Cytotoxic T cell secretion was stimulated by seeding the cells on anti-CD3ε antibody. Granules are stained through expression of pH-insensitive granzyme B-tdTomato knock-in. Movies were acquired at 10 Hz; video display speed is ×2. The exocytotic events is displayed on the left. On the right is the fluorescence intensity graph for each event aligned at the detected frame.

**Supplementary Video 3. Exemplary video of an exocytosis event of a lytic granule visualized with pH-insensitive fluorescent membrane protein.** Cytotoxic T cell secretion was stimulated by seeding the cells on anti-CD3ε antibody. Granules were labelled through expression of Synaptobrevin-mRFP knock-in. Movies were acquired at 10 Hz; video display speed is ×2. The exocytotic events is displayed on the left. On the right is the fluorescence intensity graph for each event aligned at the detected frame. Video acquired for Estl *et al*.[1].

**Supplementary Video 4. Exemplary video of an exocytosis event of a lytic granule visualized with pH-sensitive fluorescent membrane protein.** Cytotoxic T cell secretion was stimulated by seeding the cells on anti-CD3ε antibody. Granules were labelled through over-expression of Synaptobrevin-pHuji. Movies were acquired at 10 Hz; video display speed is ×2. The exocytotic events is displayed on the left. On the right is the fluorescence intensity graph for each event aligned at the detected frame.

**Supplementary Video 5. Exemplary video of individual synaptic transmission events between dorsal root ganglion neurons and spinal cord neurons.** Synaptic vesicles were labeled by over-expression of SypHy. Movies were acquired at 10 Hz, video display speed is ×2. This video displays four different regions from the same acquired movie. The left panels display four different exocytic events, up-left shows part of stimuli region, up-right shows non-sync short-BG event, down-left shows non-sync short, and down-right shows the non-sync event. The second panel shows the fluorescence intensity graph for each event aligned at the detected frame.

1. Estl, M. et al. Various Stages of Immune Synapse Formation Are Differently Dependent on the Strength of the TCR Stimulus. *Int J Mol Sci* **21** (2020).
